# Neoantigen-reactive CD8^+^ T cell engagement marks exceptional survivors of pancreatic cancer

**DOI:** 10.64898/2026.09.17.752426

**Authors:** Marc Hilmi, Joshua D. Schoenfeld, Wilson McKerrow, Yuval Elhanati, Catherine A. O’Connor, Shigeaki Umeda, Nicolas Lecomte, Jerry P. Melchor, Agustin Cardenas, Maria Lao, Vincenzo Nasca, Elias-Ramzey Karnoub, Nuray Tezcan, Zeynep Tarcan, Joanne F. Chou, Roshan Sharma, Junmin Song, Michael May, Fiyinfolu Balogun, Kenneth H. Yu, Kevin Soares, Charlotte Reiche, Martina Milighetti, Dana Pe’er, Santosha A. Vardhana, Marinela Capanu, Olca Basturk, Benoit Rousseau, Yohyoh Wang, Jayon Lihm, Neeman Mohibullah, Vineet Rolston, Ernesto Santos, Marsha Reyngold, Alice Wei, Vinod Balachandran, Mara H. Sherman, Nadeem Riaz, Christine Iacobuzio-Donahue, Benjamin Greenbaum, Eileen M. O’Reilly, Wungki Park

## Abstract

Pancreatic cancer (PC) is largely refractory to immune checkpoint blockade (ICB), although homologous recombination-deficient (HRD) tumors may derive benefit. In the POLAR trial of maintenance pembrolizumab plus olaparib after platinum-based chemotherapy for metastatic PC, responses remained heterogeneous. To define determinants of productive antitumor immunity, we integrated longitudinal blood TCR sequencing with tumor single-cell and spatial profiling. Durable benefit was associated with rare tumor-infiltrating, peripherally expanding (TIE) CD8^+^ T cell clonotypes, a subset of which were functionally neoantigen-reactive. TIE^⁺^ patients showed markedly prolonged survival beyond established genomic and immune biomarkers. Conversely, resistance was associated with spatial T cell exclusion, myCAF-rich stromal remodeling, basal-like/KRAS-associated malignant-cell programs, and expansion of CTLA4^⁺^ regulatory T cells linked to local immunosuppressive remodeling. These findings define a clonotype-resolved framework for immune monitoring and identify complementary stromal, tumor-intrinsic, and regulatory immune barriers that may guide rational combination immunotherapy in PC.

## I- Introduction

Pancreatic cancer (PC) remains one of the most lethal malignancies and is largely refractory to immune checkpoint blockade (ICB)^1,2^. This resistance has been attributed to poor immunogenicity, extensive stromal fibrosis, and a profoundly immunosuppressive tumor microenvironment^3^. As a result, immunotherapy has remained ineffective in unselected patients, underscoring the need for biologically informed strategies to identify subsets of PC capable of effective immune engagement.

Recent clinical and molecular studies have challenged the concept that PC is uniformly non-immunogenic^3^. Genomically defined subsets of PCs harbor homologous recombination deficiency (HRD), a defect in DNA damage repair that can generate high-quality mutation-derived neoantigens^4,5^. Consistent with this increased immunogenic potential, HRD tumors may derive benefit from immunotherapy-based combinations in selected clinical contexts^6-9^. However, enrichment for immunogenic mutations alone is insufficient to ensure durable antitumor immune activity and clinical response. Despite intrinsically pro-immunogenic features such as high-quality neoantigens and nucleic acid sensing^5^, effective tumor clearance mandates a permissive immune ecosystem, as resistance emerges when cell-extrinsic constraints override intrinsic immunogenicity.

The Phase II POLAR trial evaluated maintenance pembrolizumab plus olaparib in patients with metastatic PC who achieved disease control after platinum-based chemotherapy, thereby targeting a low burden disease state enriched for favorable biology and increasing the likelihood of subsequent immunologic disease control for better immune clearance^9^. Briefly, patients were prospectively stratified into three molecular cohorts based on homologous recombination repair status: cohort A, comprising tumors with pathogenic alterations in core HRD (cHRD) genes (*BRCA2, BRCA1,* or *PALB2*); cohort B, including tumors with non-core HRD (ncHRD) alterations (e.g., *RAD51B/C/D, ATM, CHEK2, BAP1, BARD1, or BRIP1*); and cohort C, consisting of patients lacking germline HR-gene mutation but platinum-sensitive HR-proficient (HRP) tumors. While clinical benefit was more pronounced in patients with cHRD, substantial inter-patient heterogeneity was observed both within the cHRD subgroup and across the broader POLAR population, highlighting that genomic stratification alone is insufficient to predict therapeutic response.

In the primary report of the POLAR trial, increased tumor-infiltrating lymphocytes were associated with clinical benefit^9^, suggesting that pre-existing immune engagement plays a critical role in mediating response. This observation raised two key questions: (1) what defines effective versus ineffective T cell responses in PC, and (2) why does only a subset of tumors translate immune infiltration into durable tumor control?

Host immune responses against cancer are dynamic processes. T cell repertoires evolve over time, clonotypes expand or contract, and functional programs of different cell compartments in the TME adapt in response to antigen exposure, stromal cues, and therapeutic pressure^10,11^. Hence, having a unique view of cellular identities and spatial organizations over longitudinal trajectories offers insight into multidimensional temporospatial dynamics. Here, we comprehensively integrate single-cell transcriptomics, spatial profiling, and longitudinal immune cell tracking from a genomically-stratified immunogenic PC dataset (POLAR trial) to define the ecosystem-level mechanisms governing immune engagement and failure in PC.

## II- Results

### 1) Clinical benefit from POLAR is associated with CD8^+^ T cell infiltration and permissive immune ecosystem

Updated survival analyses from the POLAR trial, continue to demonstrate that the clinical activity of maintenance pembrolizumab plus olaparib in patients with metastatic PC was significantly associated with HRD status (**Fig. 1A-B**). Median progression-free survival (PFS) was 8.3 months (95% CI, 5.3–39.7) in the cHRD cohort, 4.8 months (95% CI, 4.0–11.7) in the ncHRD cohort, and 3.3 months (95% CI, 1.9–4.8) in the HRP cohort. Three-year PFS rates were 30.0% (95%CI: 17.7%-50.7%), 6.7% (95%CI: 1.0%-44.3%), and 0%, respectively. With a median follow-up of 50.6 months (95% CI, 42.0–not reached [NR]), median overall survival (OS) was 27.7 months (95% CI, 11.8–NR) for cHRD, 18.2 months (95% CI, 13.0–NR) for ncHRD, and 10.4 months (95% CI, 8.9–23.8) for HRP. Three-year OS rates were 39.0% (95%CI: 24.5%-60.7%), 20.0% (95%CI: 7.3%-55.0%), and 6.7% [95%CI: 1.0%-44.3%], respectively. Despite the superior survival in patients with HRD, responses within each molecular subgroup were heterogeneous with PFS ranging from 1.35 to 65.6 months, underscoring that genomic stratification alone is insufficient to explain clinical outcome.

**Figure 1.**
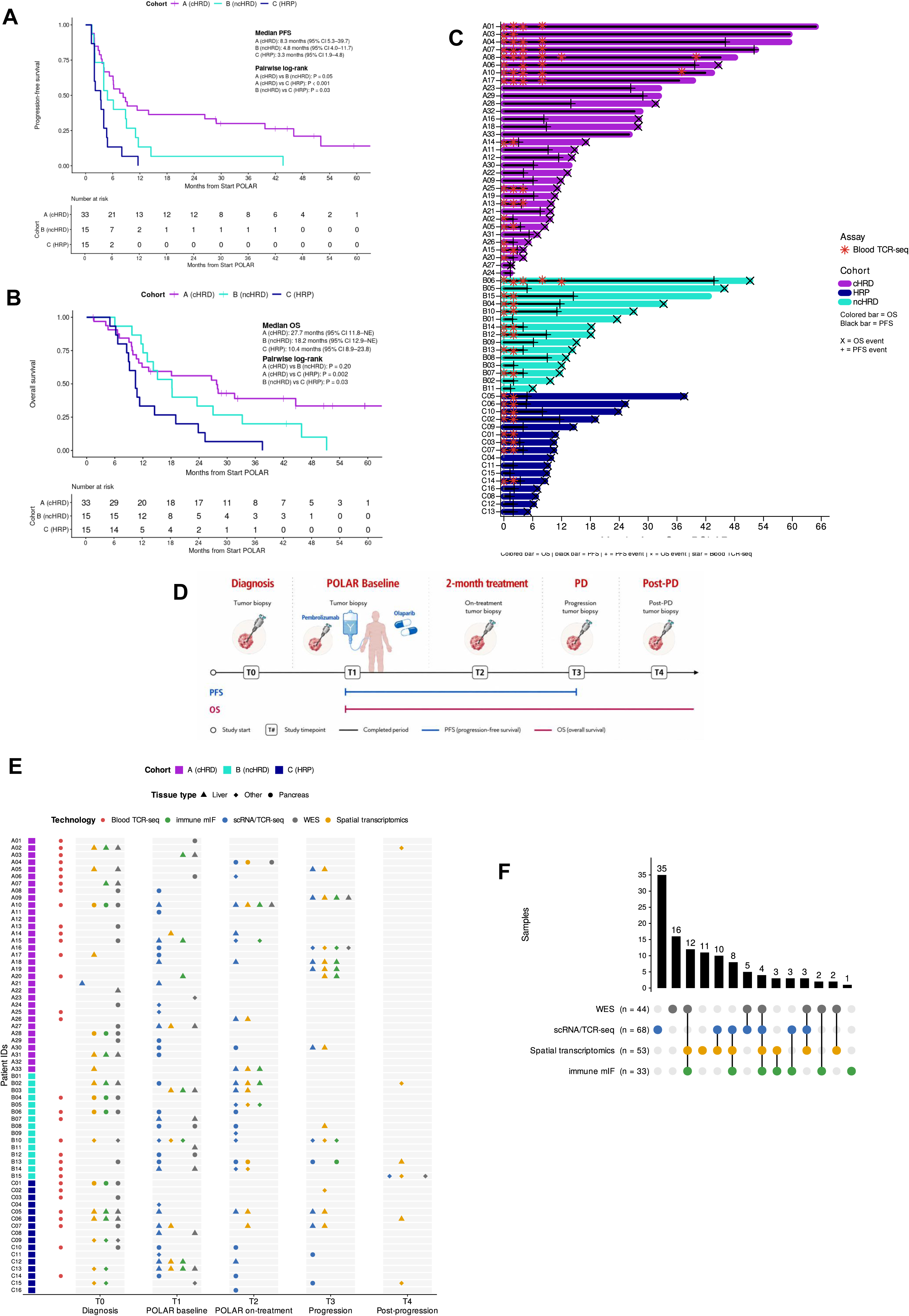
Clinical outcomes and longitudinal biospecimen profiling of the POLAR cohort. A, B, Kaplan–Meier curves for progression-free survival (PFS; A) and overall survival (OS; B) according to POLAR cohort: cohort A, core homologous recombination deficiency (cHRD); cohort B, non-core HRD (ncHRD); and cohort C, homologous recombination proficient (HRP). Median survival estimates with 95% confidence intervals and pairwise log-rank *P* values are indicated. Numbers at risk are shown below each plot. C, Swimmer plot showing individual patient PFS and OS from POLAR treatment initiation. Colored bars represent OS and are colored by cohort; overlaid white bars represent PFS. Crosses indicate PFS and OS events, and red asterisks indicate blood TCR-sequencing timepoints. D, Schematic of longitudinal sample collection during the POLAR study. T0 corresponds to diagnosis, T1 to POLAR baseline, T2 to on-treatment sampling at 2 months, T3 to progressive disease (PD), and T4 to post-progression sampling. E, Longitudinal overview of biospecimen profiling across patients and study timepoints. The left color strip indicates cohort. Red circles indicate availability of blood TCR-seq. For tissue-based assays, color denotes technology (immune mIF, scRNA/TCR-seq, WES, or spatial transcriptomics) and symbol shape denotes tissue site (liver, pancreas, or other). F, UpSet plot summarizing sample-level overlap among WES, scRNA/TCR-seq, spatial transcriptomics, and immune mIF. Bars indicate the number of samples in each exact assay combination; connected colored dots indicate the assays included in each intersection. Total numbers of profiled samples for each assay are shown alongside the matrix.

In the primary analysis of the POLAR trial, increased intratumoral CD8^+^ T cell infiltration was associated with clinical benefit^9^, suggesting that effective antitumor immunity may depend on the ability to generate and sustain tumor-directed T cell responses. We therefore performed serial peripheral blood TCR sequencing in 32 patients sampled at ≥2 timepoints to track clonal expansion and relate systemic T cell dynamics to clinical outcomes (**Fig. 1C**). To interrogate the underlying mechanisms of treatment response and resistance, longitudinal tumor biopsies were collected, when feasible, at diagnosis (T0), after induction chemotherapy and before maintenance therapy (T1; POLAR baseline), after 2 months of maintenance treatment (T2), at disease progression (T3), and post-progression (T4) (**Fig. 1D-F**). Whole-exome sequencing (WES) was performed in 42 out of the 63 patients. Paired single-cell RNA/T cell receptor (TCR) sequencing (scRNA/TCR-seq) was performed on 68 tumor samples from 45 patients, while spatial transcriptomic profiling using a fully custom 480-gene Xenium panel was conducted on 53 samples from 36 patients (**Supplementary Fig. 1**).

POLAR responders (PFS >6 months) showed greater CD8^+^ T cell infiltration than non-responders (PFS <6 months), with increased stromal CD3+CD8+ cells by multiplex immunofluorescence and higher CD8^+^ T cell abundance by spatial transcriptomics (**Supplementary Fig. 2**). Since effective anti-tumor immunity in PC depends not only on immune cell abundance, but also tissue organization, we next examined the spatial architecture of the baseline tumor microenvironment (**Extended Fig. 1A**). Niche analysis identified 12 discrete multicellular neighborhoods, one of which was preferentially enriched in responders (Niche 11) and was characterized by relative enrichment of T cell and dendritic-cell (DC) populations (**Extended Fig. 1B-D**). Within Niche 11, conventional DC (cDC) displayed increased expression of *CCR7*, a hallmark of migratory antigen-presenting states (**Extended Fig. 1E**). Moreover, cDCs localized significantly closer to tumor cells in responders than in non-responders, supporting more effective local antigen capture and presentation (mean 67 µm vs 88 µm, p=0.047, **Extended Fig. 1F**).

Acknowledging that effective T cell responses also require tumor-cell antigen presentation, we next examined whether malignant cells in responders expressed features consistent with enhanced immune recognition. Baseline malignant cells from responders expressed higher levels of antigen-presentation machinery, including TAP1, TAP2, B2M and multiple HLA class I genes (**Extended Fig. 1G**). Thus, before treatment initiation, tumors from patients who subsequently had long-term benefit from POLAR were already characterized by a permissive immune ecosystem combining CD8^+^ T cell infiltration, migratory DC niches, and preserved tumor antigen presentation (**Extended Fig. 1H**).

### 2) Treatment-associated CD8^+^ T cell clonal expansion identifies long-term responders

Given the association between CD8^+^ T cell infiltration and clinical benefit, we next asked whether specific CD8^+^ T cell states were associated with response. scRNA-seq identified 10 intratumoral CD8^+^ T cell states (**Fig. 2A, Supplementary Fig. 3**). The cluster of proliferating CD8^+^ T cells was enriched in tumors prior to progression (T1 & T2) in responders compared with non-responders (p<0.05, **Fig. 2B**).

**Figure 2.**
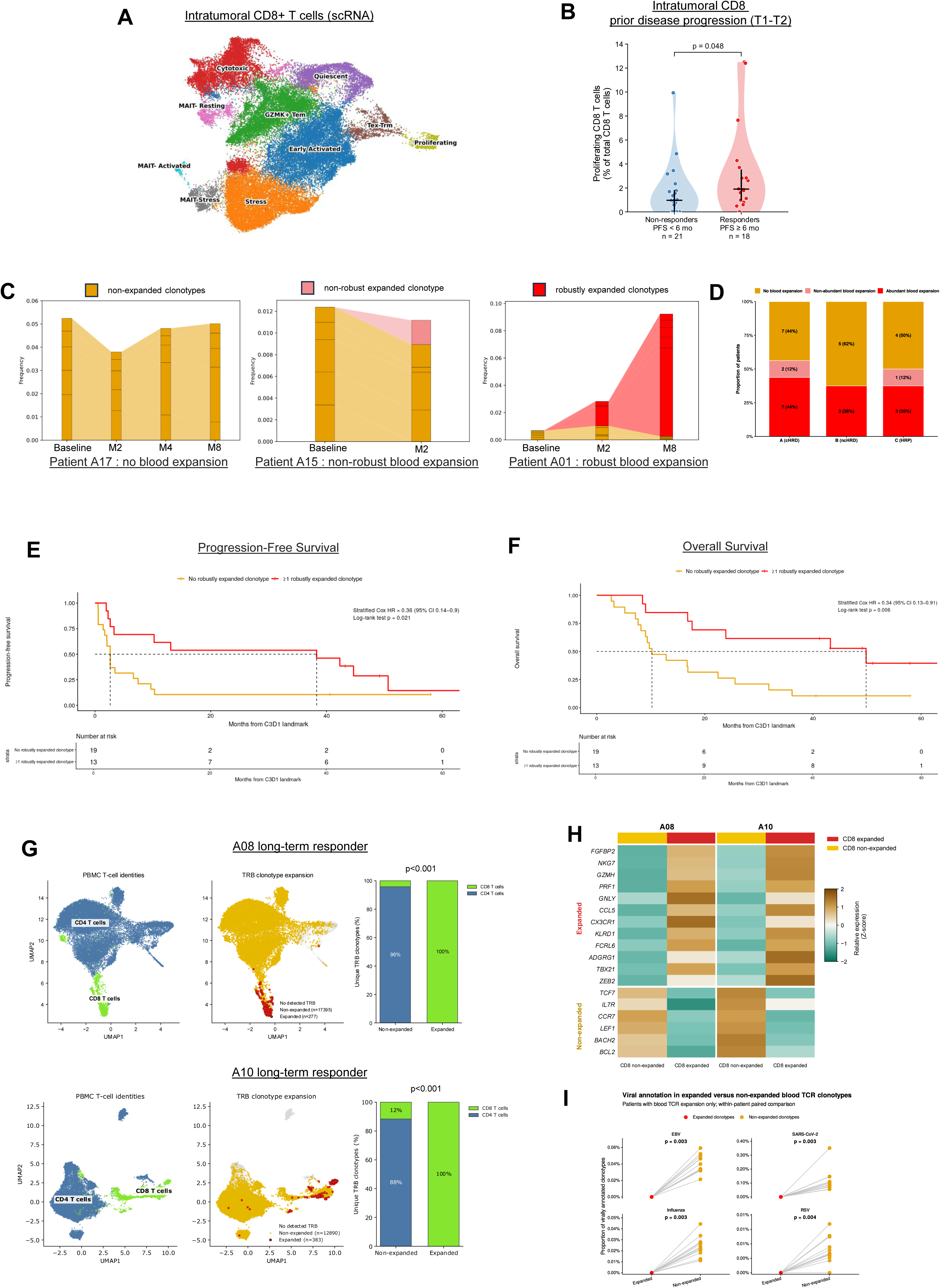
Coordinated intratumoral CD8^+^ T cell proliferation and peripheral clonotype expansion characterize clinical benefit. A, UMAP representation of intratumoral CD8^+^ T cells profiled by single-cell RNA sequencing (scRNA-seq), colored according to transcriptionally defined CD8^+^ T cell states, including cytotoxic, quiescent, GZMK^+^ effector-memory (Tem), early activated, stress, Tex-Trm, proliferating, and MAIT cell states. B, Proportion of proliferating CD8^+^ T cells among total CD8^+^ T cells in patients classified as non-responders or responders according to progression-free survival. T1 and T2 samples were pooled at the patient level before calculating cell-state proportions. Each point represents one patient. Violin plots show the distribution of values; black horizontal lines indicate the median and vertical black lines the interquartile range. Groups were compared using a two-sided Mann–Whitney U test. C, Representative longitudinal peripheral-blood TCR clonotype dynamics in patients with no detectable expansion, non-robust expansion, or robust expansion. Five non-robust clonotypes were randomly selected for visualization. Individual clonotypes are tracked across serial blood collections. D, Distribution of blood TCR expansion status across POLAR cohorts A (core homologous recombination deficiency [cHRD]), B (non-core HRD [ncHRD]) and C (homologous recombination proficient [HRP]). Patients were classified as having robust blood expansion (≥1 expanded clonotype with ≥100 TCR counts), non-robust blood expansion, or no blood expansion. Numbers and percentages of patients are indicated within bars. E,F, Kaplan–Meier estimates of progression-free survival (F) and overall survival (G) according to the presence or absence of at least one robustly expanded blood clonotype. Numbers at risk are shown below each plot; hazard ratios (HRs), 95% confidence intervals and Cox P values are indicated. G, Viral annotation of expanded and non-expanded blood TCR clonotypes for Epstein–Barr virus (EBV), SARS-CoV-2, influenza and respiratory syncytial virus (RSV). Each line connects expanded and non-expanded clonotypes from the same patient; patients without blood TCR expansion were excluded. P values for within-patient paired comparisons are shown. H, Peripheral blood mononuclear cell (PBMC) profiles from long-term responders A08 and A10. UMAPs show PBMC cell identities (left) and TRB clonotype expansion status (middle), including expanded, non-expanded and cells without detected TRB clonotypes. Stacked bars (right) show the distribution of unique expanded and non-expanded TRB clonotypes between CD8^+^ T cells and other T cells; P values are indicated. I, Heatmap of relative expression of selected effector/cytotoxic and less-differentiated T cell genes in expanded and non-expanded CD8^+^ T cells from patients A08 and A10. Expression values are scaled per gene (z-score).

This prompted us to examine whether this proliferative phenotype was accompanied by systemic T cell clonal expansion during treatment. Across all longitudinal samples, we identified 3,943,756 unique TCR sequences among 32 patients (16/32 from cohort A, 8/16 from cohort B, and 8/16 from cohort C) and only 279 (0.007%) exhibited statistically significant expansion during treatment. Marked inter-patient heterogeneity in clonal dynamics was observed, without restriction to a single molecular subgroup (p=0.89) (**Fig. 2C-D**). When all expanded clonotypes were considered, blood expansion trended toward improved survival (n=16 with blood expansion; n=16 without blood expansion), both for PFS (HR=0.53; log-rank p=0.15) and OS (HR=0.52, log-rank p=0.07; **Supplementary Fig. 4**). Because the statistical confidence of longitudinal expansion calls depends on the number of observations supporting each clonotype, we next distinguished robust blood expansion, defined by the presence of at least one expanded clonotype reaching ≥100 peripheral TCR counts. This refinement re-classified 3 patients with early progression (PFS <4 months) who had non-robust expansion. The presence of at least one robustly expanded clonotype identified a subgroup with significantly improved PFS (HR=0.36, log-rank p=0.02) and OS (HR=0.34, log-rank p=0.006; **Fig. 2E-F**).

To characterize these treatment-expanded clonotypes, we integrated blood TCR sequences with paired peripheral blood mononuclear cell (PBMC) scRNA/TCR-seq in two long-term responders (patients A08, PFS 45.0 months; patient A10, PFS 41.9 months). Expanded clonotypes were 100% CD8^+^ T cells in both patients (**Fig. 2G**) and displayed a transcriptional program distinct from non-expanded CD8^+^ clonotypes, characterized by increased expression of effector and cytotoxic genes (**Fig. 2H**). Across all patients, expanded clonotypes were patient-specific and none matched TCRs annotated as reactive to common viral antigens, including EBV, SARS-CoV-2, influenza, and RSV (**Fig. 2I**), supporting preferential expansion of non-viral CD8^+^ T cell clonotypes that may include tumor-reactive populations. Neither tumor mutational burden, nor predicted neoantigen burden was associated with the emergence of blood TCR expansion (p=0.32 and p=0.35, respectively), indicating that antigen load alone was insufficient to distinguish productive antitumor responses from other systemic T cell expansions (**Supplementary Fig. 5**).

Because T cell expansions were associated with clinical benefit, we next asked whether this signal could be captured by global repertoire-level metrics. Baseline peripheral TCR clonality did not differ between patients who subsequently developed blood expansion and those who did not (p=0.81) and remained unchanged from T1 to T2 in both groups (p=0.98 and p=1.00, respectively; **Supplementary Fig. 6**). Thus, clinically relevant T cell responses were not reflected by broad repertoire-wide oligoclonality, but rather potentially by the emergence of discrete robustly expanded clonotypes during treatment.

### 3) Pre-existing tumor-infiltrating expanded (TIE) clonotypes exhibit a cytotoxic phenotype, proximal to tumor and are associated with exceptional clinical benefit

We next asked whether treatment-expanded peripheral clonotypes infiltrated the tumor, reasoning that such blood-expanded, tumor-infiltrating clonotypes would represent candidate tumor-reactive T cell populations. Among 279 significantly expanded circulating TCR clonotypes, 27 (9.7%) were also detected in tumor tissue by scRNA/TCR-seq, thereby defining a discrete subset of TIE clonotypes (**Fig. 3A**). Notably, 21 of 27 (77.8%) TIE^⁺^ clonotypes met the robust expansion criteria, and all TIE^⁺^ patients (n=7) had robust blood expansion.

**Figure 3.**
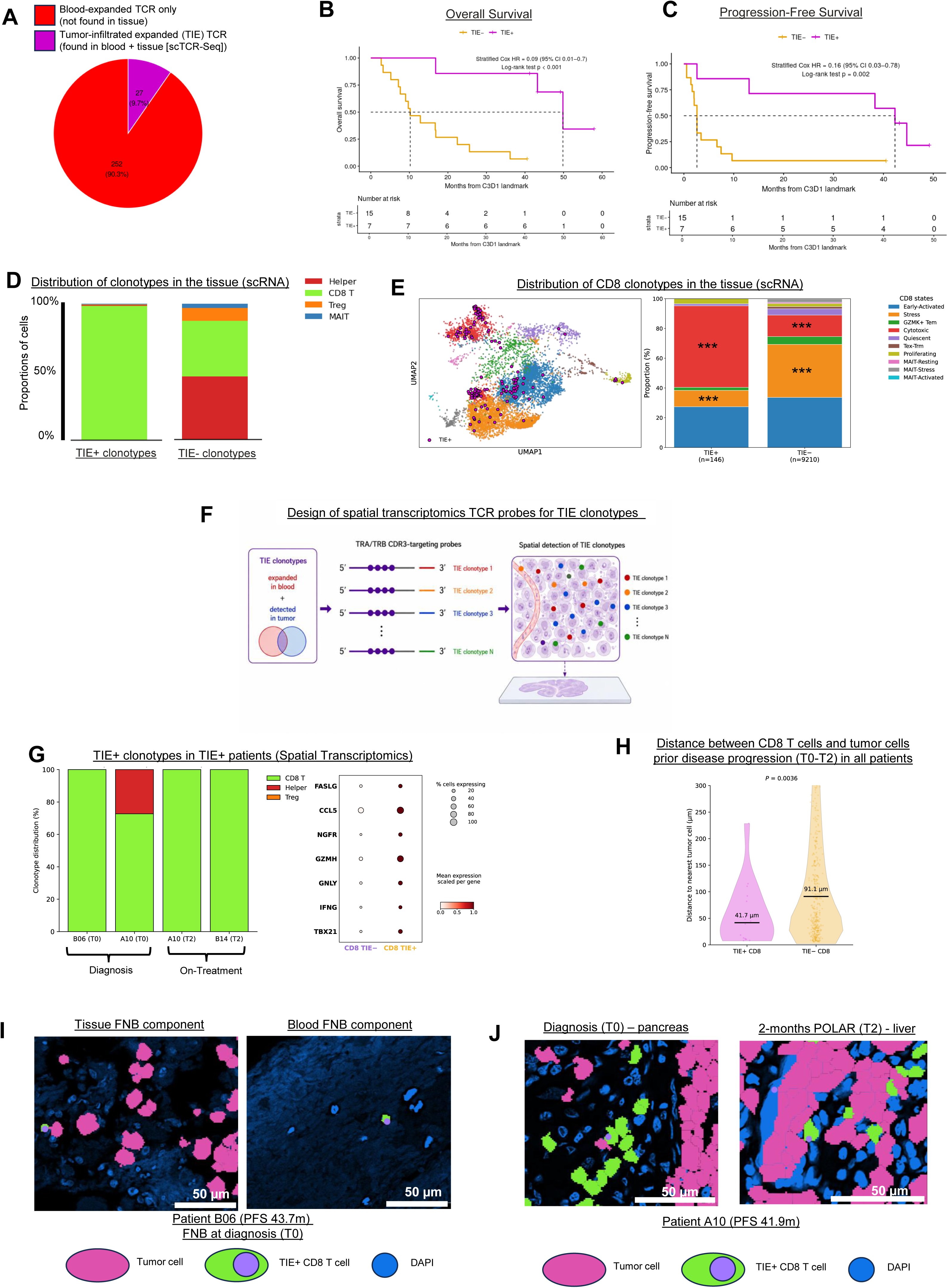
Identification and characterization of tumor-infiltrated expanded TCR clonotypes. A, Proportion of blood-expanded clonotypes identified as tumor-infiltrating expandable (TIE) clonotypes versus those not detected in the tumor. B,C, Kaplan–Meier estimates of overall survival (B) and progression-free survival (C) according to the presence (TIE+) or absence (TIE−) of TIE clonotypes. Numbers at risk are shown below each plot; hazard ratios (HRs), 95% confidence intervals and Cox P values are indicated. D, Cell-type distribution of TIE+ and TIE− clonotypes detected in tumor tissue by single-cell RNA/TCR sequencing, classified as helper T cells, CD8^+^ T cells, regulatory T cells (Tregs) or MAIT cells. E, UMAP representation of tumor CD8+ T cell states with TIE+ cells highlighted in purple (left), and relative distribution of CD8^+^ T cell states among TIE+ (n=146) and TIE− (n=9,210) cells (right). Asterisks indicate statistically significant differences between groups. F, Schematic of the strategy used to design custom spatial-transcriptomic TCR probes. Expanded clonotypes were first identified longitudinally in peripheral blood by bulk TCR sequencing, matched to tumor-infiltrating clonotypes identified by scTCR-seq/scRNA-seq, and subsequently targeted using custom Xenium TRA/TRB CDR3-specific probes for spatial localization in tumor tissue. G, Spatial-transcriptomic characterization of TIE+ clonotypes in TIE+ patients. Stacked bars show cell-type attribution of TIE+ clonotypes in samples from patients B06, A10 and B14 at diagnosis (T0) or on treatment (T2); the dot plot shows expression of selected effector/cytotoxic genes in CD8^+^ TIE− and CD8^+^ TIE+ cells. Dot size represents the percentage of cells expressing each gene and color represents mean expression scaled per gene. H, Distribution of the distance from CD8^+^ TIE^+^ and CD8^+^ TIE^-^ cells to the nearest tumor cell across pre-progression samples (T0–T2). Horizontal black lines and labels indicate medians. P value was calculated using a two-sided Mann–Whitney U test. I, Representative Xenium images of the tissue and blood components of the diagnostic fine-needle biopsy (FNB) from patient B06 J, Representative longitudinal Xenium images from patient A10, showing the pancreatic tumor at diagnosis (T0) and a liver metastasis after 2 months of POLAR treatment (T2). In I,J, tumor cells are shown in magenta, CD8^+^ T cells in green, TIE-specific TCR probe signals in purple, and nuclei (DAPI) in blue. Green CD8^+^ T cells without a purple signal represent CD8^+^ T cells in which the targeted TIE clonotype was not detected. Scale bars, 50 µm.

Despite representing a minority of expanded blood clonotypes, the presence of at least one TIE clonotype identified patients with markedly improved clinical outcomes. These patients had OS of at least 18 months despite metastatic disease, and demonstrated markedly improved OS compared to TIE^-^ patients (HR=0.09, 95% CI 0.01-0.70, log rank p<0.001; **Fig. 3B**). The 3-year OS rate was 85.7% (95% CI 63.3-100) in TIE^+^ patients compared with 13.3% (95% CI 0.04–48.4) in TIE^-^ patients. Consistent with this observation, TIE^+^ patients also experienced markedly prolonged PFS compared with TIE^-^ patients (HR=0.16, 95% CI 0.03-0.78, p=0.002; **Fig. 3C**). Median PFS was 42.2 months in TIE^⁺^ patients, whereas TIE^⁻^ patients had a median PFS of 2.5 months. TIE clonotypes were not shared across patients, and were detected across cohorts A (cHRD, n=3) and B (ncHRD, n=4), but not in cohort C (HRP), indicating that tumor infiltration by expanded clonotypes was restricted to HRD tumors only.

Among the 7 patients harboring TIE clonotypes, 4 had tumor tissue available at baseline (T1), and in all cases these clonotypes were already detectable prior to POLAR maintenance. To assess their temporal dynamics, we performed extended longitudinal blood TCR sequencing in TIE^⁺^ patients, profiling at least five timepoints from baseline to >3 years on POLAR treatment. TIE clonotypes showed durable persistence, with 12 of 14 evaluable clonotypes (85.7%) remaining detectable beyond 3 years across three TIE^⁺^ patients (**Supplementary Fig. 7**), demonstrating that these clonotypes can persist long-term. Sequential tumor samples at T1 and T2 were available for 3 TIE^+^ patients. The frequency of these shared clonotypes did not significantly increase at T2 compared with T1 (p>0.05), arguing against intratumoral clonal expansion. This observation is consistent with prior reports that pre-existing intratumoral T cell clones have limited reinvigoration capacity following PD-1 blockade, supporting the notion that productive antitumor responses may depend on expansion outside the tumor at draining lymph nodes and subsequent tumor infiltration^12^.

Consistent with the blood findings, TIE^+^ TCR clonotypes were predominantly found within CD8^+^ T cells (143/145, 98.6%) (**Fig. 3D**), in contrast to TIE^-^ TCR clonotypes detected in tumor tissue without evidence of blood expansion, which showed more heterogeneous cellular T cell subsets (p<0.001). We next examined the intratumoral state of these TIE^⁺^ CD8^+^ T cells. TIE^⁺^ cells were strongly enriched for the cytotoxic CD8^+^ state compared with TIE^⁻^ CD8^+^ T cells (54.8% vs 14.4%; p<0.001), whereas the stress-associated state was depleted (11.0% vs 35.7%; p<0.001; **Fig. 3E**). Within the cytotoxic CD8^+^ T cell state, TIE^⁺^ cells expressed significantly higher levels of effector and activation genes, including *IFNG, TNF, TNFSF9, CD83, PRF1*, and *NKG7*, than TIE^⁻^ cells (**Supplementary Fig. 8**). Finally, TIE^+^ CD8^+^ T cells showed increased clonotype convergence (i.e. presence of distinct T cell clones that share identical or highly similar CDR3 amino acid sequences despite arising from different nucleotide rearrangements) relative to TIE^-^ CD8^+^ T cells in TIE^+^ patients (48% versus 24%; p<0.001), further supporting a shared antigen-driven effector program^13^ (**Supplementary Fig. 9**).

We then sought to spatially localize these clonotypes directly in tissue. To this end, we designed custom Xenium probes targeting TCR CDR3 sequences from TIE^+^ clonotypes, enabling their spatial visualization in tumor sections (**Fig. 3F**). Spatial profiling was conducted on 6 tumor samples from 5 of 7 TIE^⁺^ patients with available tissue, including 2 treatment-naïve samples collected at diagnosis (T0), 3 on-treatment (T2) and 1 at progression (T3). TIE clonotypes were detected in five of six samples, including both treatment-naïve tumors, demonstrating that these clonotypes were already present within the tumor before exposure to POLAR therapy. The single negative sample was profiled using only one short TIE TCR sequence, potentially limiting detection sensitivity. Consistent with scRNA/TCR-seq, spatially detected TIE clonotypes localized predominantly to CD8^+^ T cells and displayed increased expression of cytotoxic and effector genes compared with TIE^⁻^ CD8^+^ T cells (**Fig. 3G**). Prior to disease progression, TIE^+^ CD8^+^ T cells tended to localize closer to tumor cells than TIE^-^ CD8^+^ T cells, consistent with more intimate tumor interactions (42 µm vs 91 µm; p=0.004; **Fig. 3H**). In fine-needle biopsy material from patient B06 (PFS 43.7 months), we could capture both a tissue component and a blood-associated component, thereby allowing direct visualization of circulating and stromal compartments within the same specimen (**Fig. 3I**). In a second example, TIE^+^ CD8^+^ T cells were detected across longitudinal tumor samples from patient A10 (PFS 41.9 months), including a pancreatic biopsy obtained at diagnosis and a liver sample collected on treatment, demonstrating persistence of TIE^⁺^ clonotypes across timepoints and anatomical sites (**Fig. 3J**).

Collectively, these analyses suggest that TIE clonotypes are not merely defined by their presence in both blood and tumor, but correspond to a distinct biological state characterized by CD8^+^ T cell lineage restriction, neoantigen-cognate T cell clonal expansion, and proximity to tumor cells. These properties distinguish TIE^⁺^ clonotypes from other expanded T cell populations and provide a potential mechanistic basis for their strong association with exceptional clinical benefit observed in select patients on the POLAR trial.

### 4) TIE^+^ clonotypes include functionally validated neoantigen-reactive T cells

Having established that TIE clonotypes exhibit features consistent with antigen-driven antitumor immunity, we next asked whether they directly recognized tumor-derived neoantigens. We focused on two long-term responders, A08 and A10, for whom PBMC scRNA/TCR-seq (**Fig. 2G**) and tumor scRNA/TCR-seq were available. Following WES-based neoantigen quality prediction using *Neoqual*, we synthesized peptides spanning the top 10 predicted MHC class I neoepitopes, together with the corresponding mutant KRAS peptide, and stimulated autologous PBMCs in vitro for 14 days, as previously described^11^. After peptide restimulation, CD8^+^ T cells were sorted into CD107a^⁺^ and CD107a^⁻^ fractions and subjected to bulk TCRβ sequencing (**Fig. 4A**). TCRβ sequences recovered from these functional fractions were matched to paired TCRα/TCRβ clonotypes identified by PBMC scTCR-seq and subsequently to tumor scTCR-seq, allowing neoantigen-responsive cells to be directly linked to previously defined TIE clonotypes.

**Figure 4.**
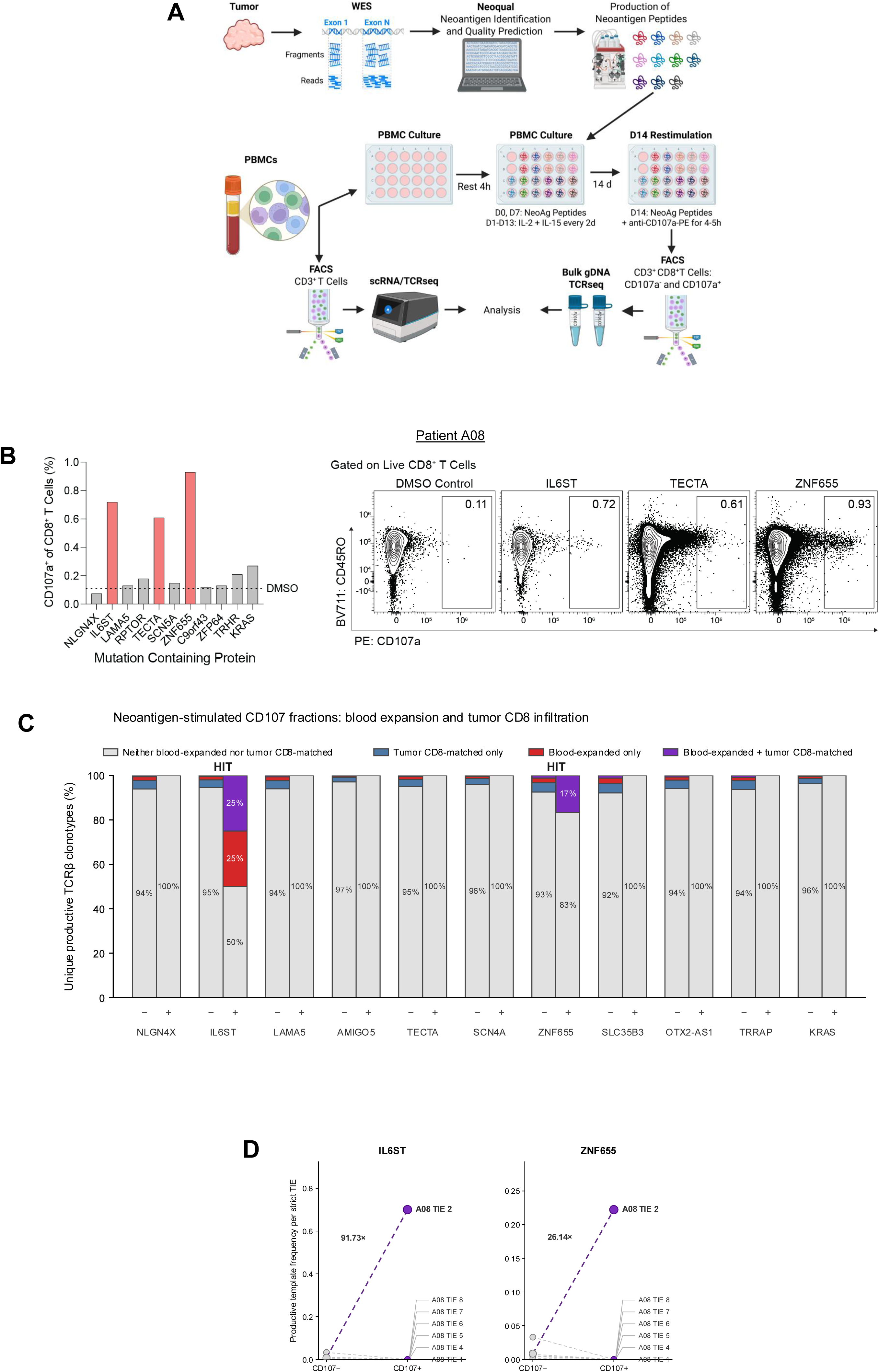
Functional neoantigen validation and TCR clonotype tracking of neoantigen-reactive CD8^⁺^ T cells. A, Schematic of the neoantigen-specific T cell functional validation workflow. Tumor whole-exome sequencing (WES) was used to identify candidate neoantigens and generate patient-specific neoantigen peptides. PBMCs were cultured with neoantigen peptides and IL-2/IL-15 for 14 days, followed by peptide restimulation and CD107a-based sorting of CD3^⁺^CD8^⁺^ T cells. In parallel, baseline PBMC CD3^⁺^ T cells underwent single-cell RNA/TCR sequencing. Bulk TCR sequencing of CD107a^⁺^ and CD107a^⁻^ fractions was integrated with single-cell TCR data to identify and functionally characterize neoantigen-reactive T cell clonotypes. B, Functional neoantigen screen in patient A08. Left, percentage of CD107a^⁺^ cells among CD8^⁺^ T cells following stimulation with peptides containing the indicated tumor mutations; the dotted line indicates the DMSO control. Right, representative flow-cytometry plots showing CD107a expression after DMSO, IL6ST, TECTA and ZNF655 peptide stimulation. Numbers indicate the percentage of CD107a^⁺^ cells within live CD8^⁺^ T cells. C, TCRβ clonotype composition of the CD107a^⁻^ and CD107a^⁺^ fractions following stimulation with the indicated A08 neoantigen peptides. Bars show the proportion of unique productive TCRβ clonotypes classified according to their detection among blood-expanded clonotypes and/or matched tumor CD8^⁺^ T cell clonotypes: neither (grey), tumor CD8^+^-matched only (blue), blood-expanded only (red), or both blood-expanded and tumor CD8^+^-matched (purple). Percentages are calculated within each sorted fraction. HIT denotes peptide-stimulated CD107a^⁺^ fractions enriched in clonotypes that were blood-expanded and tumor CD8^+^-matched. D, Productive template frequencies of individual tumor-infiltrating expanded (TIE) clonotypes are shown in CD107a^−^ and CD107a^+^ fractions following stimulation with IL6ST (left) or ZNF655 (right). Each dashed line connects the frequency of the same TIE clonotype between the CD107a^−^ and CD107a^+^ fractions. Grey circles indicate CD107a^−^ frequencies and purple circles indicate CD107a^+^ frequencies. A08 TIE 2 is highlighted in purple; fold changes between CD107a^−^ and CD107a^+^ frequencies are indicated.

In patient A08, peptides derived from TECTA, ZNF655, and IL6ST induced the strongest CD107a responses above the DMSO control, corresponding to 5.5-, 8.5-, and 6.5-fold increases, respectively (**Fig. 4B**). TCR sequencing of the corresponding CD107a fractions identified selective enrichment of A08 TIE-2 following IL6ST and ZNF655 stimulation (**Fig. 4C-D**). Thus, a clonotype previously shown to expand in blood and infiltrate the tumor exhibited functional reactivity against patient-specific neoantigens. IL6ST encodes gp130, a central co-receptor of the IL-6 signaling pathway, which plays a key role in PC progression^14^, whereas ZNF655 has been reported to be overexpressed in PC and to promote tumor progression through cell cycle regulation^15^.

We independently observed a similar pattern in patient A10, in whom 5 of 10 predicted neoantigen peptides induced CD107a responses above DMSO, including peptides derived from AATK, SCN4A, ARL11, SYT16, and NR1H4 (**Supplementary Fig. 10A**). Among the three A10 TIE clonotypes, TIE-1 was selectively detected in the CD107a^⁺^ fraction following SYT16 stimulation, whereas TIE-2 was detected across multiple peptide conditions and TIE-3 was not recovered (Supplementary Fig. 10B). A10 TIE-1 was enriched 2.13-fold in the SYT16-stimulated CD107a^⁺^ relative to CD107a^⁻^ fraction (**Supplementary Fig. 10C**), providing a second example of functional neoantigen recognition by a TIE clonotype.

We next confirmed that these candidate neoantigens were supported by both genomic and transcriptional evidence within the tumor. The IL6ST and ZNF655 mutations in A08 had variant allele frequencies (VAFs) of 0.12 and 0.28, respectively, while the SYT16 mutation in A10 had a VAF of 0.24. Moreover, all three corresponding source genes were detected in malignant cells at POLAR baseline, compared with only 5 of 12 source genes for non-reactive candidate peptides (**Supplementary Fig. 11**).

Together, these findings suggest that TIE clonotypes can include functionally neoantigen-reactive CD8^+^ T cells targeting patient-specific tumor mutations expressed by the malignant compartment.

### 5) Stromal exclusion, basal-like tumor state, and KRAS signaling accompany resistance

We next investigated mechanisms limiting productive T cell-tumor interactions and associated with resistance. In TIE^⁺^ patients, the median distance between TIE^⁺^ CD8^+^ T cells and malignant cells increased from 42 µm prior to progression to 82 µm at progression (p=0.08), consistent with acquired spatial exclusion (**Fig. 5A-B**). Similarly, at POLAR baseline, CD8^+^ T cells were located significantly farther from malignant cells in non-responders than in responders (111 µm vs 73 µm, p<0.001; **Fig. 5C**).

**Figure 5.**
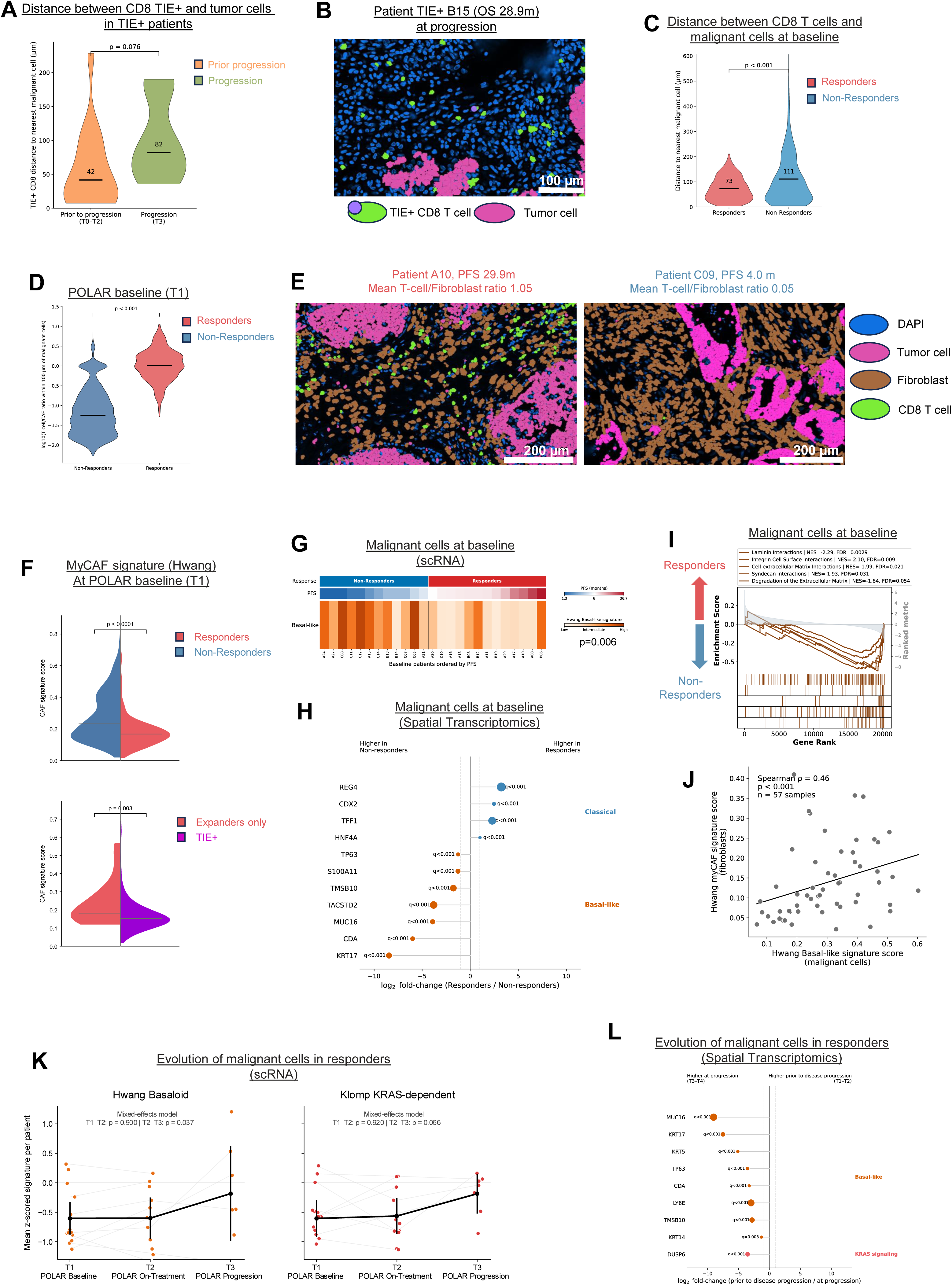
T cell exclusion, myofibroblastic stroma, basal-like tumor states, and KRAS signaling are associated with disease progression. A, Distance from TIE^+^ CD8^+^ T cells to the nearest malignant cell in TIE^+^ patients before disease progression (T0–T2) and at progression (T3). Horizontal lines indicate medians (42 and 82 μm, respectively); two-sided P value is shown. B, Representative spatial image from TIE^+^ patient B15 (OS, 28.9 months) at progression showing tumor cells (pink), TIE^+^ CD8^+^ T cells (green with TIE signal indicated in purple), and DAPI (blue). Scale bar, 100 μm. C, Distance from CD8^+^ T cells to the nearest malignant cell at POLAR baseline (T1) in responders (PFS ≥6 months) and non-responders (PFS <6 months). Horizontal lines indicate medians (73 and 111 μm, respectively); P value is shown. D, Distribution of the log10-transformed T cell/CAF ratio within 100 μm of malignant cells at baseline in responders and non-responders. Horizontal lines indicate medians; P value is shown. E, Representative spatial images from responder A10 (PFS, 29.9 months; mean T cell/fibroblast ratio, 1.05) and non-responder C09 (PFS, 4.0 months; mean T cell/fibroblast ratio, 0.05), showing fibroblasts (gray), tumor cells (pink), and CD8+ T cells (green). Scale bars, 200 μm. F, Hwang myofibroblastic signature scores in fibroblasts at POLAR baseline. Top, comparison between responders and non-responders; bottom, comparison between patients with blood-expanded clonotypes without detected tumor infiltration (“Expanders only”) and patients with tumor-infiltrating and blood-expanded clonotypes (TIE+). Horizontal lines indicate medians; Mann–Whitney P values are shown. G, Hwang Basal-like signature in malignant cells by scRNA-seq at baseline. Patients are ordered by PFS and annotated by response status and PFS; the P value for the responder versus non-responder comparison is shown. H, Differential expression of selected Classical (blue) and Basal-like (orange) genes in malignant cells profiled by spatial transcriptomics at baseline. Points indicate log2 fold-change in mean normalized expression (responders/non-responders), with positive values indicating higher expression in responders and negative values higher expression in non-responders. Benjamini–Hochberg-adjusted q values are shown. I, Gene-set enrichment analysis of baseline malignant-cell transcriptional profiles comparing responders and non-responders. Enrichment curves are shown for extracellular-matrix-associated pathways, with normalized enrichment scores (NES) and FDR values indicated in the legend. J, Association between the Hwang myCAF signature score in fibroblasts and the Hwang Basal-like signature score in malignant cells across 57 samples. The fitted linear trend is shown. K, Longitudinal evolution of malignant-cell transcriptional states by scRNA-seq among responders. Mean patient-level z-scored Hwang Basaloid (left) and Klomp KRAS-dependent (right) signature scores are shown at POLAR baseline (T1), on-treatment (T2), and progression (T3). Gray lines connect longitudinal samples from individual patients and black lines summarize the group trajectory. P values for T1–T2 and T2–T3 comparisons from mixed-effects models are shown. L, Gene-level changes in malignant cells profiled by spatial transcriptomics among responders, comparing samples prior to disease progression (T1–T2) with samples at progression (T3–T4). Basal-like genes showing greater than two-fold change toward progression and DUSP6, representing KRAS signaling, are displayed. Points indicate log2 fold-change (prior to progression/at progression); negative values indicate higher expression at progression and positive values higher expression before progression. Dashed lines denote two-fold changes (log2 fold-change ±1), and Benjamini–Hochberg-adjusted q values are shown.

Analysis of response-associated spatial niches identified Niche 0, a fibroblast-rich neighborhood preferentially enriched in non-responders (**Extended Fig. 1B, Extended Fig. 2A-B**). Consistent with a stromal barrier to T cell access, the local T cell-to-fibroblast ratio around malignant cells was markedly lower in non-responders (p<0.001; **Fig. 5D-E**). Spatial differential expression analysis of fibroblasts within Niche 0 revealed significant enrichment of extracellular-matrix and myofibroblastic genes, including *COL11A1*, *COL12A1*, *POSTN*, *COL1A1*, *MMP11*, *VCAN*, and *TAGLN* (**Extended Fig. 2C**). Independently, scRNA-seq confirmed enrichment of myCAF-associated genes and higher myCAF signature scores in fibroblasts from non-responders at baseline (**Fig. 5F, Extended Fig. 2D**). Notably, among patients with blood T-cell expansion, myCAF scores were higher in those whose expanded clonotypes failed to infiltrate the tumor than in TIE^⁺^ patients (**Fig. 5F**), consistent with myofibroblastic stroma limiting tumor access by expanded circulating T cells. Histologically, this phenotype was reflected by the emergence of a poorly cellular, deserted stromal architecture after induction chemotherapy: among 11 patients with paired H&E samples at diagnosis (T0) and POLAR baseline (T1), 7 (63.6%) developed deserted stroma, which was associated with trends toward shorter PFS (HR=3.39, p=0.13) and OS (HR=3.88, p=0.10; **Extended Fig. 2E-H**).

We next examined whether malignant-cell states were coupled to this resistant stromal phenotype. At baseline, malignant cells from non-responders showed increased basal-like transcriptional programs by both scRNA-seq and spatial transcriptomics (**Fig. 5G-H**), together with enrichment of pathways related to laminin, integrin, and extracellular-matrix interactions (**Fig. 5I**). Across samples, malignant-cell basal-like scores positively correlated with fibroblast myCAF scores (ρ=0.46, p<0.001; **Fig. 5J**), supporting coordinated tumor-stromal states associated with primary resistance. Finally, we asked whether similar malignant programs emerged during acquired resistance. Among responders with longitudinal samples, basal-like programs increased at progression, accompanied by increased KRAS-dependent signaling (**Fig. 5K-L**).

Overall, these data support a model in which myofibroblastic stromal remodeling and basal-like tumor programs cooperate to restrict T cell access and promote both primary and acquired resistance.

### 6) CTLA-4-associated immunosuppression emerges during resistance

Compensatory CTLA-4 upregulation, particularly within Tregs, has been implicated in resistance to PD-1 blockade in different solid tumors ^16-18^. We therefore asked whether CTLA-4-associated immunosuppression emerged during resistance to pembrolizumab plus olaparib.

Among 8 patients with paired scRNA-seq samples, the proportion of *CTLA4*^⁺^ Tregs significantly increased at progression compared with prior timepoints (p=0.031; **Fig. 6A**). This increase was independently observed by spatial transcriptomics in 3 patients with paired samples (p<0.001; **Fig. 6A**). Pathway analysis of Tregs at progression further revealed enrichment of immunosuppressive programs, including IL-10, IL-35, and IL-2 family signaling, together with proliferative programs involving DNA replication and chromosome segregation (**Fig. 6B**).

**Figure 6.**
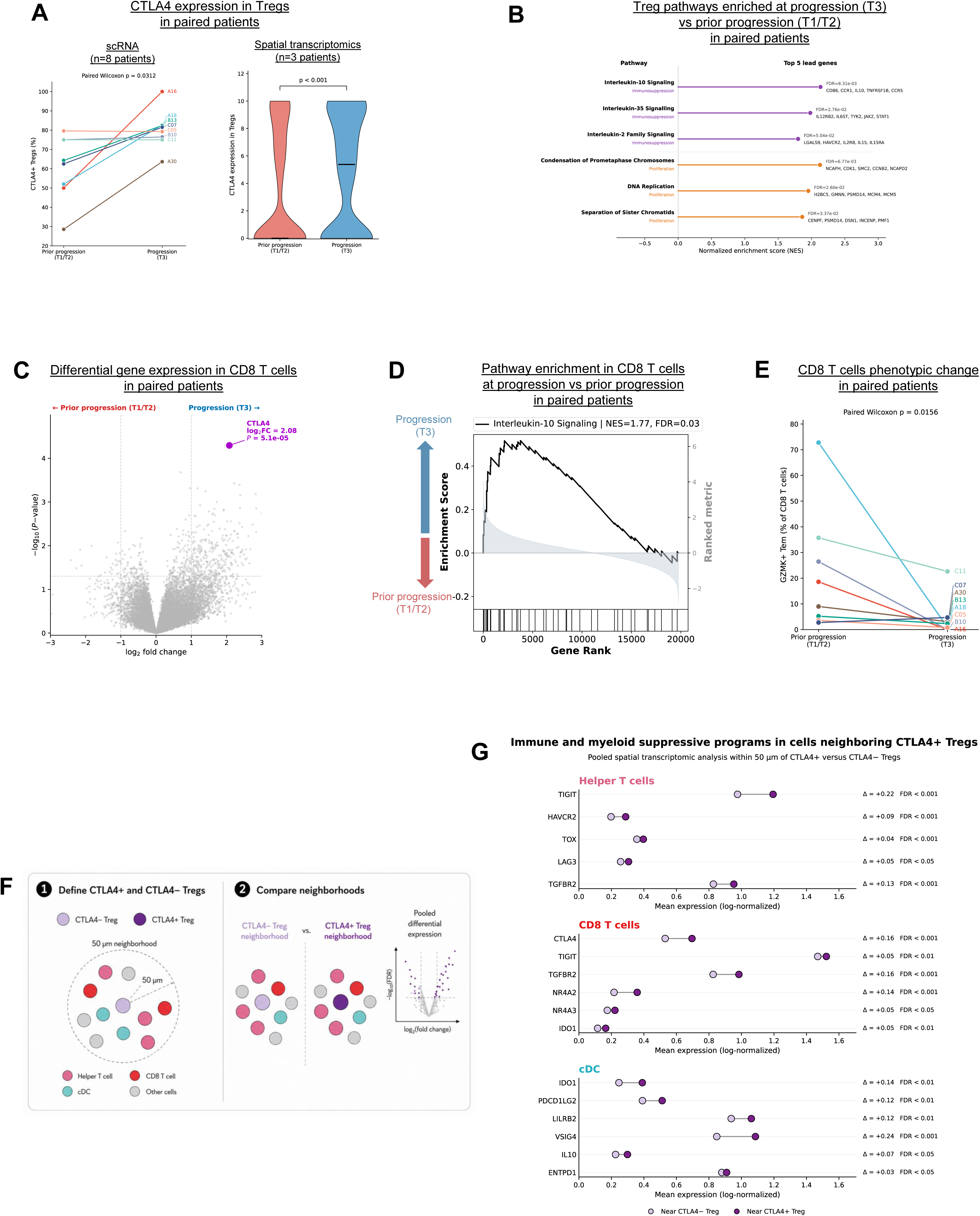
CTLA4 increases at progression and defines a spatially immunosuppressive immune niche. A, CTLA4 expression in Tregs in patients with paired pre-progression (T1/T2) and progression (T3) samples. Left, paired analysis of the proportion of CTLA4^⁺^ Tregs by scRNA-seq in eight patients before disease progression (T1–T2) and at progression (T3). Lines connect measurements from individual patients; significance was assessed using a two-sided paired Wilcoxon test. Right, CTLA4 expression in Tregs profiled by spatial transcriptomics in three paired patients before progression (T1–T2) and at progression (T3). Horizontal lines indicate medians; two-sided Mann–Whitney P value is shown. B, Gene-set enrichment analysis of Tregs at progression versus pre-progression, highlighting enrichment of immunoregulatory pathways, together with proliferative programs. NES, normalized enrichment score. C, Differential gene-expression analysis of CD8^+^ T cells in paired scRNA-seq samples at progression versus pre-progression. CTLA4 is highlighted among top gene increased at progression. D, GSEA demonstrating enrichment of IL-10 signaling in CD8+ T cells at progression. E, Paired analysis of the proportion of GZMK^+^ effector-memory CD8^+^ T cells before and at progression; lines connect samples from the same patient. F, Schematic of the spatial-neighborhood analysis. Tregs were classified as CTLA4+ or CTLA4− based on CTLA4 detection, and 50-µm neighborhoods were defined around each population. Helper T cells, CD8+ T cells and cDCs located exclusively within CTLA4+ or CTLA4− Treg neighborhoods were compared; cells falling within both neighborhoods were excluded. Differential expression analysis was performed after pooling cells across evaluable tumors. G, Spatial transcriptional programs in immune cells neighboring CTLA4^⁺^ Tregs. Mean expression of selected immunoregulatory genes in Helper T cells (top), CD8+ T cells (middle), and cDCs (bottom) located within 50 μm of CTLA4^⁺^ versus CTLA4^⁻^ Tregs across spatially profiled tumor samples. Light and dark purple points indicate cells neighboring CTLA4^⁻^ and CTLA4^⁺^ Tregs, respectively, and connecting lines indicate the difference in mean log-normalized expression. Δ denotes the difference in mean expression between CTLA4^⁺^- and CTLA4^⁻^-Treg neighborhoods (CTLA4^⁺^ minus CTLA4^⁻^). Statistical significance was assessed using two-sided Mann–Whitney tests with Benjamini–Hochberg correction for multiple testing; FDR thresholds are indicated.

Concomitant changes were observed in the CD8^+^ T cell compartment. *CTLA4* was the most strongly upregulated genes at progression (**Fig. 6C**), accompanied by enrichment of IL-10 signaling (**Fig. 6D**). In parallel, the proportion of GZMK^⁺^ T effector memory (Tem) cells significantly decreased at progression in paired patients (p=0.016; **Fig. 6E),** consistent with loss of this pre-existing effector-memory state.

We next asked whether *CTLA4*^⁺^ Tregs were associated with localized immunosuppressive remodeling. Using spatial transcriptomics, we compared cells within 50 µm of *CTLA4*^⁺^ versus *CTLA4*^⁻^ Tregs (**Fig. 6F**). Helper T cells neighboring *CTLA4*^⁺^ Tregs showed increased expression of inhibitory and dysfunctional programs, including *TIGIT, HAVCR2, TOX*, and *LAG3*, while neighboring CD8^+^ T cells showed increased *CTLA4*, *TIGIT*, *TGFBR2*, and *NR4A2/3*. cDCs within these neighborhoods similarly expressed higher levels of regulatory myeloid genes, including *IDO1, PDCD1LG2, LILRB2*, and *VSIG4* (**Fig. 6G**). These findings identify *CTLA4*^⁺^ Treg expansion as a feature of acquired resistance associated with coordinated local immunosuppressive remodeling across T cell and myeloid compartments.

## III- Discussion

Pancreatic cancer (PC) remains largely refractory to ICB because of poor tumor immunogenicity compounded by stromal restriction and immune suppression^3^. Even within HRD PC, benefit remains heterogeneous^9^, indicating that genomic immunogenicity alone is insufficient. By integrating longitudinal blood TCR sequencing with paired tumor scRNA/TCR-seq, spatial transcriptomics, and histopathology in POLAR, we identify TIE^+^ CD8^+^ T clonotypes as a rare but biologically meaningful correlate of exceptional survivors of metastatic PC treated with pembrolizumab and olaparib.

Clinical benefit was not fully explained by HRD status, CD8^+^ T cell abundance, peripheral expansion, TMB, neoantigen burden, or global TCR clonality alone. Instead, TIE clonotypes represented a minor subset of treatment-expanded clones that were detectable before POLAR therapy in evaluable untreated tumors, persisted longitudinally, localized near malignant cells, adopted cytotoxic programs, and in selected patients, directly recognized patient-specific neoantigens. This extends observations that effective immunotherapies and neoantigen vaccines can amplify limited antigen-specific T cell populations^10,11,19-21^. TIE clonotypes were observed in both cHRD and ncHRD cohorts, arguing against restriction to cHRD biology. Their absence in cohort C should be interpreted cautiously given fewer durable responses and limited sample size, rather than as evidence that TIE biology is intrinsically HRD-specific.

Our data also support convergent mechanisms of immune failure. Primary resistance was associated with myCAF-rich stroma, increased CD8^+^ T cell-to-tumor distance, and coordinated basal-like tumor states, consistent with stromal restriction of effective immune trafficking in PC^22-24^. At progression, these features were accompanied by increased KRAS-dependent signaling and greater spatial exclusion of TIE^+^ cells. Acquired resistance was additionally marked by expansion of CTLA4^+^ Tregs, *CTLA4* upregulation in CD8+ T cells, IL-10-associated programs, and localized suppressive remodeling of T cell and myeloid compartments, consistent with compensatory immune checkpoint activation after PD-1 blockade^16-18^.

These findings nominate complementary therapeutic strategies. First, amplifying tumor-reactive clonotypes through personalized neoantigen vaccines or TCR-based approaches could expand endogenous TIE-like populations or directly exploit tumor-reactive receptors identified from responding patients^10,11,21^. Second, concurrent K/RAS inhibitions^25-28^ may suppress malignant programs emerging with acquired resistance while promoting a more permissive immune ecosystem, providing a rationale for combination with immune activation. Third, remodeling myCAF/ECM programs may be required to convert systemic T cell expansion into physical tumor access^22-24^. Finally, CTLA4 blockade could relieve the regulatory state emerging at progression and provides a rationale for PD-1/CTLA4 combinations in selected HRD PC^16-18,29,30^. Thus, durable immunotherapy benefit in PC may require coordinated antigenic amplification, stromal remodeling, oncogenic pathway suppression, and effective engagement of adaptive immunity.

This study also underscores the value of prospective longitudinal biospecimen collection in PC, where limited tissue availability, low tumor cellularity, fibrosis, and challenging biopsy sites constrain translational investigation. Integrating serial blood with scarce tumor specimens across orthogonal platforms can reveal treatment-emergent biology that baseline profiling alone cannot capture.

Limitations include the small numbers of TIE^+^ patients, incomplete longitudinal tissue sampling, and analysis within a single clinical trial. Independent validation in additional PC cohorts is needed, as is determining whether TIE-like clonotypes generalize to productive antitumor immunity beyond PC. Nevertheless, the convergence of longitudinal blood and tumor TCR profiling with single-cell, spatial, functional and histologic analyses provides orthogonal evidence for a model in which systemic T cell expansion alone is insufficient for durable tumor control. Instead, long-term survival is associated with the ability of rare, neoantigen-reactive CD8^⁺^ T cell clonotypes to access and engage malignant cells within a permissive tumor ecosystem. These findings suggest that future effective immunotherapy in pancreatic cancer may require both reinvigoration of tumor-reactive T cells and optimization of the tumor ecosystem to permit their access and antitumor activity.

## Figure legends

**Extended Data Fig. 1.**
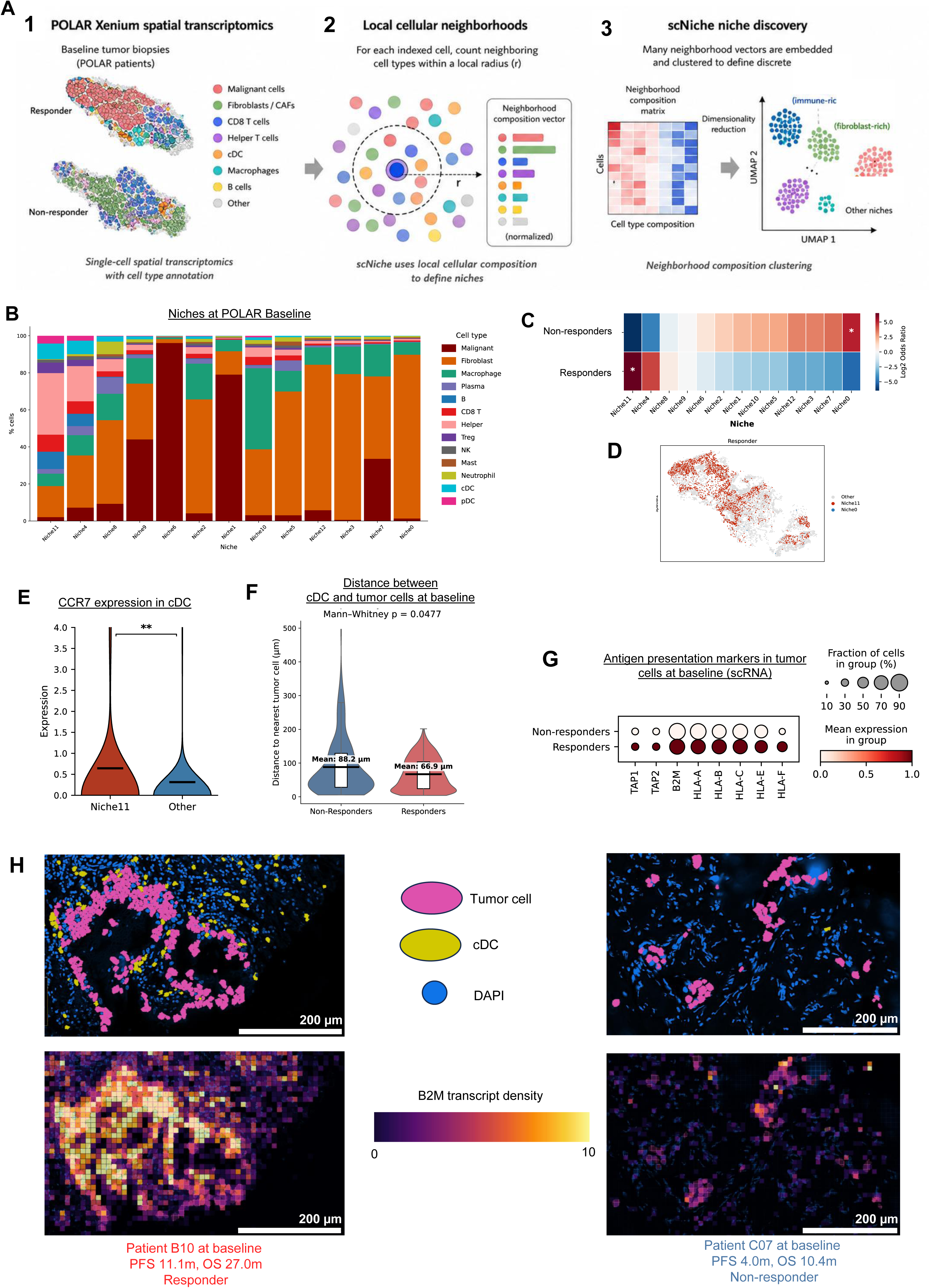
A responder-enriched spatial niche is associated with CCR7^⁺^ cDC–tumor proximity and enhanced antigen presentation at baseline. A, Schematic overview of scNiche analysis applied to POLAR Xenium spatial transcriptomic data. Baseline tumor biopsies were profiled by Xenium and annotated at single-cell resolution. For each indexed cell, the local cellular neighborhood was defined by counting neighboring cell types within a fixed spatial radius and encoding these counts as a normalized neighborhood composition vector. Neighborhood vectors were then embedded and clustered to define discrete spatial niches. B, Cell-type composition of spatial niches identified at POLAR baseline by Xenium spatial transcriptomics. Bars show the percentage of each annotated cell type within each niche. C, Association of baseline spatial niches with clinical response. Heatmap shows log2 odds ratios for enrichment of each niche in responders and non-responders. Asterisks indicate statistically enriched niches. D, Representative spatial map from a responder at POLAR baseline showing the distribution of Niche 11 and Niche 0 cells. Other cells are shown in grey. E, CCR7 expression in conventional dendritic cells located within Niche 11 compared with conventional dendritic cells located in other niches, assessed by spatial transcriptomics. Statistical significance is indicated. F, Distance from conventional dendritic cells to the nearest tumor cell at POLAR baseline in non-responders and responders. Mean distances and Mann–Whitney test P value are shown. G Dot plot showing expression of antigen-presentation genes in baseline malignant cells from responders and non-responders by scRNA-seq. Dot size indicates the fraction of cells expressing each gene, and color indicates mean expression within each group. H, Representative Xenium images from patient B10 and patient C07 at POLAR baseline. Upper panels show tumor cells, conventional dendritic cells and DAPI. Lower panels show B2M transcript density in the corresponding tissue regions. Scale bars, 200 μm.

**Extended Data Fig. 2.**
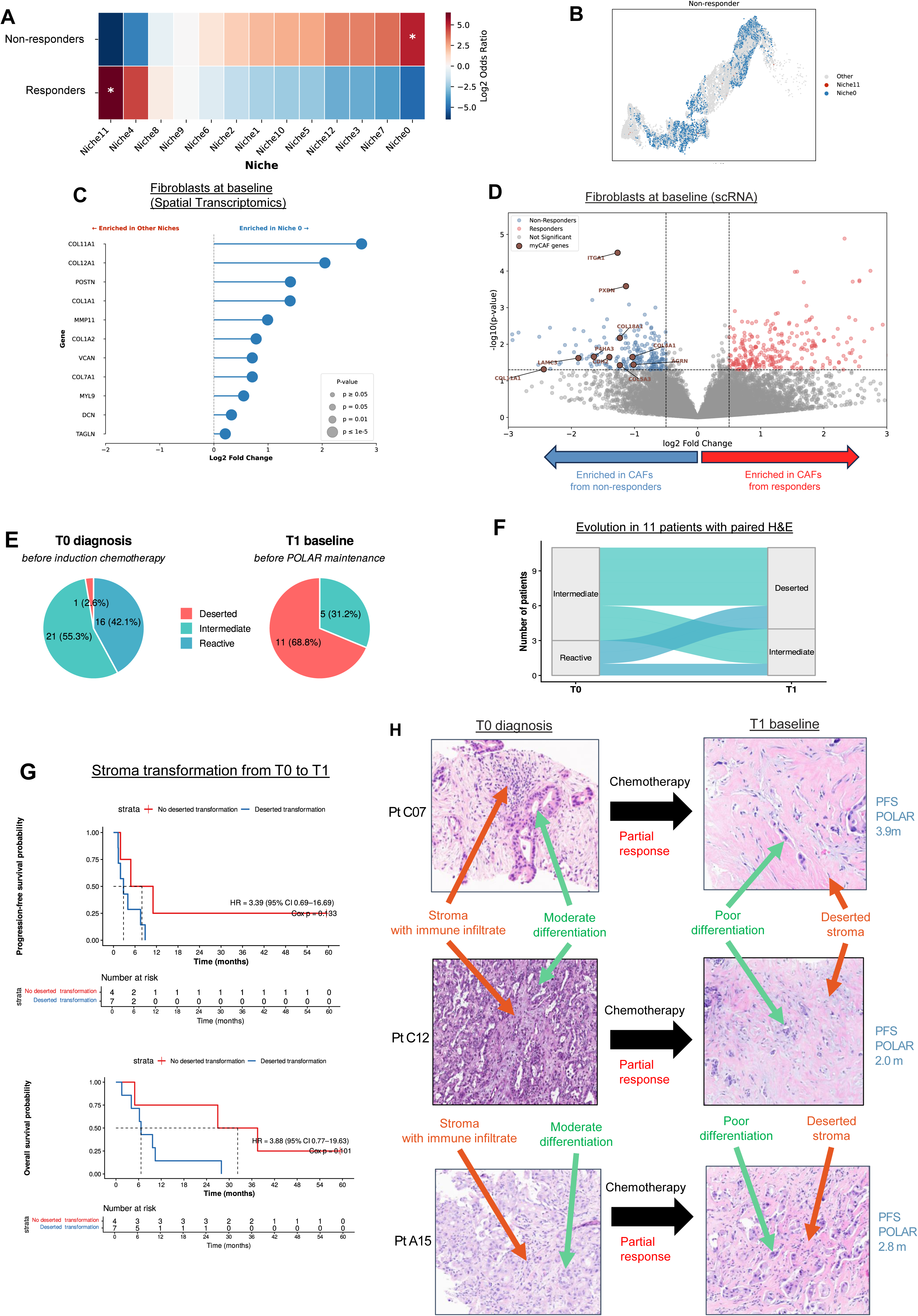
Stromal remodeling from diagnosis to POLAR baseline. A, Niche enrichment in responders and non-responders at POLAR baseline. Heatmap shows the log2 odds ratio for enrichment of each spatial niche according to response group; asterisks indicate statistically significant enrichment. B, Representative spatial map from a non-responder showing the distribution of Niche 0 and Niche 11 relative to all other cells. C, Differential gene expression in fibroblasts located within Niche 0 versus fibroblasts in other spatial niches at POLAR baseline, assessed by spatial transcriptomics. The x-axis indicates log2 fold change; point size reflects statistical significance. D, Volcano plot of differential gene expression in fibroblasts from responders versus non-responders at POLAR baseline using scRNA-seq. Genes enriched in fibroblasts from non-responders and responders are shown in blue and red, respectively; selected myofibroblastic CAF-associated genes are annotated. E, Distribution of stromal histologic phenotypes at diagnosis (T0, before induction chemotherapy) and at POLAR baseline (T1, before POLAR maintenance). Stromal phenotypes were classified as deserted, intermediate, or reactive. Numbers and percentages of evaluable samples are shown. F, Alluvial plot showing changes in stromal phenotype from T0 to T1 in the 11 patients with paired H&E specimens. G, Kaplan–Meier estimates of progression-free survival (top) and overall survival (bottom) according to stromal transformation between T0 and T1. Patients with non-deserted stroma at T0 were classified as having a *deserted transformation* if the T1 specimen was deserted, or *no deserted transformation* if the T1 specimen remained non-deserted. Hazard ratios with 95% confidence intervals and Cox proportional-hazards P values are shown; numbers at risk are indicated below each plot. H, Representative paired H&E sections from three patients (C07, C12, and A15) at T0 and T1, illustrating changes in stromal morphology between diagnosis and POLAR baseline. Arrows indicate representative areas of immune-infiltrated or deserted stroma and differences in tumor differentiation. Best response to induction chemotherapy and PFS during POLAR maintenance are indicated for each patient.

**Supplementary Fig. 1 | Cell type annotation of single-cell and spatial transcriptomic datasets.**

(A) Uniform Manifold Approximation and Projection (UMAP) embedding of all cells profiled by single-cell RNA sequencing across the POLAR cohort. Cells are colored and annotated according to their assigned cell type identity. Major immune, stromal, and epithelial compartments are indicated.

(B) Dot plot showing expression of canonical marker genes used for cell type annotation in the single-cell RNA sequencing dataset. Dot size indicates the fraction of cells expressing a given gene within each cell type, and color intensity indicates average normalized expression.

(C) UMAP embedding of all cells profiled by spatial transcriptomics across the POLAR cohort. Cells are colored and annotated according to their assigned cell type identity. Major immune, stromal, endothelial, and epithelial populations are indicated.

(D) Dot plot showing expression of canonical marker genes used for cell type annotation in the spatial transcriptomic dataset. Dot size indicates the fraction of cells expressing a given gene within each cell type, and color intensity indicates average normalized expression.

**Supplementary Figure 2 | CD8 T-cell abundance and transcriptional states according to clinical response.**

A, Quantification of CD3^⁺^CD8^⁺^ cells among stromal cells by multiplex immunofluorescence in POLAR, comparing non-responders and responders. Points are colored by molecular cohort. P value was calculated using a two-sided Mann–Whitney test.

B, Percentage of CD8 T cells among immune cells measured by Xenium spatial transcriptomics in POLAR, comparing non-responders and responders. Points are colored by molecular cohort. P value was calculated using a two-sided Mann–Whitney test.

C, Representative Xenium images from a non-responder and a responder at POLAR baseline. Tumor cells, CD8 T cells and DAPI are shown as indicated. Scale bars, 500 μm.

**Supplementary Figure 3 | CD8**^⁺^ **T-cell clustering and metabolic pathway scores.**

A, Dot plot showing the expression of selected marker genes across CD8^⁺^ T-cell clusters identified by single-cell RNA sequencing. Dot size represents the fraction of cells expressing each gene within a cluster, and color intensity represents the relative mean expression scaled from 0 to 1 for each gene. Marker genes are grouped according to the indicated transcriptional programs.

B, Violin plots comparing the Quiescent cluster with all other CD8^⁺^ T-cell clusters. Violins show cell-level score distributions; internal box plots indicate the median and interquartile range, with whiskers extending to 1.5 times the interquartile range. Statistical significance was assessed using two-sided Mann–Whitney U tests with Benjamini–Hochberg correction across the six comparisons. ****q < 0.0001.

**Supplementary Fig. 4 | Progression-free and overall survival according to peripheral blood TCR expansion.**

A, Kaplan–Meier estimates of progression-free survival according to the presence or absence of at least one significantly expanded blood TCR clonotype. Hazard ratio, 95% confidence interval, Cox model P value, and numbers at risk are shown.

B, Kaplan–Meier estimates of overall survival according to the presence or absence of at least one significantly expanded blood TCR clonotype. Hazard ratio, 95% confidence interval, Cox model P value, and numbers at risk are shown.

**Supplementary Fig. 5 | Neoantigen burden and tumor mutational burden according to blood TCR expansion.**

A, Predicted neoantigen burden in patients with or without blood TCR expansion. Each point represents one patient. P value is shown.

B, Tumor mutational burden, expressed as mutations per megabase, in patients with or without blood TCR expansion. Each point represents one patient. P value is shown.

**Supplementary Figure 6 | Peripheral TCRβ clonality at baseline and on treatment according to clonal expansion status.**

A, Blood TRB clonality at POLAR baseline (T1) in patients with versus without subsequent blood TCR expansion. Each point represents one patient; the P value is shown.

B, Paired blood TRB clonality at POLAR baseline (T1) and on-treatment (T2), shown separately for patients without blood TCR expansion and patients with blood TCR expansion. Lines connect measurements from the same patient; paired P values are shown.

Supplementary Figure 7 | Longitudinal tracking of tumor-infiltrating expanded (TIE) clonotypes in peripheral blood.

Peripheral frequencies of individual TIE clonotypes are shown longitudinally for each TIE^⁺^ patient from baseline through long-term follow-up. Each line represents a distinct clonotype, with different shades used to distinguish clonotypes within each patient. Filled circles indicate detected clonotypes, open circles indicate clonotypes that were assayed but not detected (frequency = 0), and crosses indicate time points for which no sample was collected or available. Time points correspond to baseline, months 2 (M2), 4 (M4), 8 (M8), and 12 (M12), 12–36 months (M12–36), and >36 months (>M36).

## Methods

### Trial design, patients and endpoints

POLAR was a single-institution, open-label, non-randomized phase 2 trial conducted at Memorial Sloan Kettering Cancer Center (NCT04666740). Participants with metastatic PC, aged ≥18 years and with ECOG performance status 0–1, were enrolled into three biomarker-defined cohorts: Cohort A: tumors harboring core-HRD (cHRD) mutations (*BRCA1/2* or *PALB2),* Cohort B: tumors harboring non-core HRD (ncHRD) gene mutations (*ATM, BAP1, BARD1, BLM, BRIP1, CHEK2, FAM175A, FANCA, FANCC, NBN, RAD50, RAD51, RTEL1, MUTYH*), Cohort C: homologous-recombination proficient (HRP) tumors with durable platinum sensitivity.

Eligible participants had histologically confirmed pancreatic carcinoma (including ductal adenocarcinoma, adenosquamous carcinoma or acinar cell carcinoma), completed platinum-based induction therapy, and had no disease progression prior to enrollment. Cohort A and B for 4 months and Cohort C for 6 months. Exclusion criteria included >2 prior lines of systemic therapy, prior PARP inhibitor (PARPi) or immune checkpoint blockade (ICB), or radiographic progression before trial entry.

Participants received maintenance pembrolizumab (anti–PD-1 antibody) and olaparib (PARPi). All participants received maintenance olaparib (300 mg BID) and pembrolizumab (every 3 weeks for 6 months, then every 6 weeks until disease progression or unacceptable toxicity). Select patients were allowed to continue beyond radiographic progression if deriving clinical benefit. Enrollment occurred from December 28, 2020, to February 20, 2024 (data cutoff: June 25, 2026).

We conducted the study in accordance with the Declaration of Helsinki and good clinical practice guidelines. The study was approved by the institutional review board (IRB) at Memorial Sloan Kettering Cancer Center (MSK) and the United States Federal Drug Administration (FDA). All participants provided written informed consent.

### Survival analyses

Median follow-up was estimated using the reverse Kaplan–Meier method. OS and PFS were calculated from the date of treatment initiation to death (OS) and to the first occurrence of disease progression or death (PFS), respectively. OS and PFS were estimated using the Kaplan–Meier method and reported separately for the three POLAR cohorts.

For analyses of longitudinal blood TCR metrics, including clonal expansion, robust clonal expansion, and tumor-infiltrating expanded (TIE) clonotypes, landmark analyses were performed at cycle 3 day 1 (C3D1; week 6), corresponding to the first on-treatment blood assessment. Only patients alive and progression-free at the landmark were included in PFS analyses, and only patients alive at the landmark were included in OS analyses; survival times were calculated from the landmark. Associations between blood TCR metrics and OS or PFS were evaluated using Cox proportional hazards models stratified by POLAR cohort to account for differences across study subpopulations. As a post-hoc exploratory analysis, Kaplan–Meier curves were compared using unstratified log-rank tests.

### Blood TCR Sequencing

After PicoGreen quantification and quality control by Agilent BioAnalyzer, TCR expression was profiled as described in Shugay, et al.^31^ with modifications. Briefly, 0.4-700 ng total RNA were used as input into a cDNA synthesis reaction using SMARTScribe Reverse Transcriptase (Takara catalog # 639537). The reaction was primed from TCRα and TCRβ using RT primers containing unique molecular identifiers (UMIs) and a custom 5’ Smart NNNa adapter was used. RNasin (Promega catalog # N2611) was added to the reaction, and Smart NNNNa was degraded by the addition of 20 units uracil DNA glycosylase (NEB catalog # M0280L) and incubation at 37°C for 40 minutes. First strand cDNA was then amplified with Q5 High-Fidelity DNA Polymerase (NEB catalog # M049L) for 23 cycles of PCR using UMI-containing primers specific to TCRα and TCRβ. The resulting material was split into separate α chain and β chain reactions to undergo sample barcoding with a second round of PCR amplification for 14 cycles. Purified PCR product underwent sequencing library preparation with the KAPA Hyper Prep Kit (Roche catalog # 07962363001) with 8 cycles of PCR. Samples were sequenced on the NovaSeq X in a PE150 run using the NovaSeq X 25B Reagent Kit (Illumina) for an average of 19 million paired reads per sample.

To focus on biologically meaningful systemic T cell responses and minimize stochastic fluctuations due to low-frequency clones, TCR sequences were considered expanded if the change between baseline and any subsequent timepoint was found to be statistically significantly more than 2-fold. Statistical significance was assessed using the Fisher’s exact test to compare baseline counts to half the counts at the subsequent timepoint, as previously described^10^. Baseline was defined as the earliest timepoint where at least 1000 TCRs were sequenced (i.e. for patients with poor coverage (≤1000 total TCRs) in the screening sample, we also considered expanding clones that achieved statistical significance at later timepoints). To ensure that statistically significant longitudinal expansion calls were supported by a sufficient number of observations and were less sensitive to stochastic fluctuations among low-count clonotypes, expanded clonotypes were further classified as robust if they reached ≥100 TCR counts at any sampled timepoint. TRB clonality was calculated as one minus the normalized Shannon entropy of the TRB clone-size distribution, where (f(i)) denotes the fraction of T cell receptors corresponding to sequence (i), and (n) represents the number of unique TRB sequences detected.

Viral-associated TCRB sequences were identified computationally as previously described^32^. Briefly, more than 30,000 sequenced TCRB repertoires were analyzed to identify co-occurring clusters of T cells in donors sharing the same HLA. These HLA-associated T cell clusters were grouped based on their co-occurrence with other T cell clusters associated with different HLAs, forming groups termed exposure co-occurrence clusters (ECOclusters). ECOclusters were subsequently validated by quantifying their presence and absence in repertoires with known serostatus. Clusters that are significantly enriched in repertoires from individuals positive for a given exposure, compared with those that are negative for that exposure, are labeled exposure associated.

### In vitro peptide stimulation

To identify neoantigen specific T cells, we stimulated autologous PBMCs with neoantigen-derived peptides as previously described. Specifically, tumor biopsies were analyzed by WES to identify somatic mutations which were subsequently filtered to remove low confidence mutations ^20^. MSK NeoQual™ is used to evaluate neoantigen quality based on the neoantigen quality model described by Łuksza et al. (2022). Briefly, neoantigen quality is determined by three main components: (a) the difference in MHC-binding affinity between the wild-type and mutant peptides, (b) the predicted antigenic distance between the wild-type and mutant peptides, reflecting their potential for differential T-cell recognition, and (c) the sequence similarity of the mutant peptide to known immunogenic epitopes in the Immune Epitope Database (IEDB). Higher neoantigen quality scores indicate stronger preferential presentation of the mutant peptide, greater predicted discrimination of the mutant peptide from its wild-type counterpart by T cells, and greater similarity to known immunogenic epitopes. We computationally created 27mers with the mutated residue at position 14 and synthesized four 14-15mer peptides spanning the mutated residue (residue 1-15, 5-20, 9-24, and 13-27) for each of the top 10 highest binding neoantigens and the patient-specific mutant KRAS allele. All peptides were resuspended in DMSO at 5-10□mg/mL. Cryopreserved PBMCs were thawed, added dropwise to PBMC media (RPMI-1640 + 10% fetal calf serum (FCS), 2 mM glutamine, 1X NEAA, 50 µM 2-mercaptoethanol, and 1X penicillin-streptomycin) and centrifuged at 350 x g for 7 min at 4C to remove cryopreservation media. Cells were resuspended in PBMC media, counted using trypan blue, and resuspended at 1 x 106 viable cells/mL in PBMC media. PBMCs were plated in a tissue-culture treated 48-well plate (1 mL/well; 1-2 wells per peptide or DMSO control) and allowed to rest for 3-4 h at 21% O2 in a humidity-controlled environment (37°C, 5% CO2; NuAire). In the interim, the remaining cells were subjected to fluorescence-activated cell sorting (FACS) to isolate viable CD3+ T cells for single cell gene expression and V(D)J profiling (10x Genomics), see “FACS” and “scRNAseq” sections below.

Following 3-4 h of rest, 1□×□ 106 PBMCs in a 48-well plate were stimulated with peptide pools (5 µg/mL per peptide, total peptide concentration of 20□μg/ml), and IL-2 (100□U/ml) and IL-15 (10□ng/mL) added the next day (day 1) and every subsequent 2–3□days. On day 7, PBMCs were restimulated with peptide pools. On day 14, PBMCs were restimulated with peptide pools in the presence of anti-CD107a-PE (1:50, BD Biosciences, H4A3, #555801, RRID:AB_396135) as a marker of CD8+ T cell degranulation. Following 4-5 hours of re-stimulation, PBMCs were collected and processed for FACS as below. Isolated viable CD8++ CD107a-and CD107a+ T cells were centrifuged at 700 g x 2 min, genomic DNA was extracted according to the manufacturer’s instructions (NEB Monarch® gDNA Extraction Kit), and processed for bulk TCR sequencing (Adaptive Biotechnologies, see below).

### FACS

Briefly, Single cell suspensions were centrifuged at 350 x g at 4C for 5 min and resuspended in Ghost DyeTM Viability Dye 780 (Tonbo, 1:1500) and Fc Block (Biolegend, 1:20) for 10 minutes in PBS on ice. Cells were washed in FACS buffer (PBS + 2% FCS), centrifuged at 350 x g at 4C for 5 min and resuspended cell surface staining solution composed of cell surface marker antibodies at prespecified dilutions in 1:1 BD HorizonTM Brilliant Stain Buffer (BD Biosciences) and FACS buffer. Cells were stained for 30 minutes on ice, washed in FACS buffer, and centrifuged at 350 x g at 4C for 5 min. Stained cells were resuspended in FACS buffer, filtered, and CD3+ T cells (scRNA/TCRseq) or CD3+ CD8++ CD107a- or CD107a+ (bulk TCRseq) were sorted into FACS buffer using a 70 μm nozzle on either a FACSAriaTM III or FACSymphonyTM S6 (BD Biosciences) flow cytometer in the MSK Flow Cytometry Core Facility.

FACS surface antibodies: CD8+a-BUV395 (BD Biosciences, RPA-T8, #563795, RRID:AB_2722501), CD45-BV570 (BioLegend, HI30, #304033, RRID:AB_10899568), CD3-PerCP/Cy5.5 (BioLegend, SK7, #344808, RRID:AB_10640736), CD45RO-BV711 (BD Biosciences, UCHL1, #563723, RRID:AB_2744413), HLA-DR-BV786 (BD Biosciences, G46-6, #564041, RRID: AB_2738559), CD107a-PE (BD Biosciences, H4A3, #555801, RRID:AB_396135), CD11b-PE/Cy7 (BioLegend, M1/70, #101216, RRID:AB_312799), CD19-PE/Cy7 (BioLegend, SJ25C1, #363012, RRID:AB_2564203), and CD56-PE/Cy7 (BioLegend, 5.1H11, #362510, RRID:AB_2563927).

### Histological analysis

Formalin-fixed paraffin-embedded (FFPE) tumor samples from the POLAR trial were sectioned and stained with hematoxylin and eosin (H&E) according to standard clinical pathology protocols. Only samples containing both viable tumor epithelium and assessable stromal compartments were included in the analysis.

H&E slides were independently reviewed by an expert gastrointestinal pathologist, blinded to clinical outcomes and molecular data. Tumors were classified based on stromal immune architecture according to previously published criteria^33^, into three categories:

- Deserted stroma, characterized by dense fibrotic stroma with minimal inflammatory infiltrate and sparse immune cells in proximity to tumor glands.
- Intermediate stroma, displaying focal or heterogeneous immune cell infiltration within the stromal compartment, without diffuse immune-rich regions.
- Reactive stroma, defined by prominent immune cell infiltration, including lymphoid aggregates, with active stromal remodeling and close spatial association between immune cells and tumor epithelium.

Classification was based on the dominant stromal pattern observed across the tumor section. In cases with heterogeneous features, the category representing the majority of the stromal area was assigned. Discordant assessments were resolved by joint review and consensus.

Stromal immune phenotypes were assessed at diagnosis (T0, prior to induction chemotherapy) and at baseline before POLAR maintenance therapy (T1), enabling longitudinal comparison of stromal remodeling and immune exclusion patterns.

### Single cell RNA/TCR Sequencing for tumor samples

Single-cell RNA-seq and TCR-seq libraries were generated using the Chromium GEM-X Single Cell 5′ v3 platform (10x Genomics) according to the manufacturer’s instructions. Cell concentration and viability were assessed prior to loading, and approximately 20,000 cells were targeted for recovery per sample. Cells were partitioned into Gel Bead-in-Emulsions (GEMs), followed by reverse transcription and cell barcoding. GEMs were subsequently recovered, emulsions were broken, and barcoded cDNA was purified and amplified according to the manufacturer’s protocol. Amplified cDNA was used for both 5′ gene expression (GEX) library construction and T cell receptor (TCR) enrichment. GEX libraries were generated by fragmentation, end repair and A-tailing, adaptor ligation, sample indexing PCR, and SPRIselect bead purification. For TCR-seq library construction, TCR transcripts were enriched from amplified cDNA using the human T cell V(D)J enrichment workflow (10x Genomics), followed by library construction and sample indexing according to the manufacturer’s instructions. Final GEX and TCR V(D)J libraries were sequenced on an Illumina NovaSeq X Plus using paired-end, dual-index sequencing. Raw sequencing data were processed using Cell Ranger v10.0.0 (10x Genomics). GEX reads were aligned to the 10x Genomics GRCh38-2024-A human transcriptome reference to generate gene expression count matrices. TCR V(D)J reads were processed using the Cell Ranger V(D)J pipeline for TCR sequence reconstruction and clonotype assignment.

Single-cell profiling was performed on 68 tumor samples from 45 patients enrolled in POLAR, including samples obtained at diagnosis (T0, n = 1), baseline before POLAR maintenance therapy (T1, n = 31), on treatment at approximately 2 months (T2, n = 23), and disease progression (T3, n = 13). One baseline scRNA/TCR-seq sample was obtained from patient C04, who subsequently failed screening and did not initiate POLAR treatment; this patient was therefore excluded from all treatment-response and survival analyses. Samples were derived predominantly from liver (n = 30) and pancreatic (n = 26) lesions, with additional peritoneal (n = 6) and other metastatic sites (n = 5). Gene-expression matrices were processed on a per-sample basis using Scanpy. Ambient RNA contamination was reduced using CellBender, and quality-control metrics were calculated using sc.pp.calculate_qc_metrics. Cells with fewer than 20 detected genes or >30% mitochondrial transcripts were excluded. Potential doublets were identified using Scrublet. After quality control, approximately 270,000 cells were retained for downstream analyses, with a median of approximately 4,000 cells per sample.

Gene-expression counts were normalized by library size and log-transformed. Highly variable genes were selected using the Seurat v3 method (n = 4,000), followed by principal-component analysis, construction of a k-nearest-neighbor graph, UMAP dimensionality reduction, and clustering using PhenoGraph with Leiden community detection.

Multicellular niches were inferred using scNiche^34^, which assigns each cell to a niche based on the composition of its local cellular neighborhood. To assess whether specific niches were associated with patient response, each cell was labeled according to its patient’s response status (Responder vs Non-responder, defined by PFS being higher or lower than 6 months). For each niche, we quantified enrichment using a two-sided Fisher’s exact test on a 2×2 contingency table comparing counts of Responder vs Non-responder cells inside the niche versus outside the niche. P values were adjusted across niches using Benjamini–Hochberg FDR, and effect sizes were reported as log2 odds ratios (with a 0.5 pseudocount for stability) and visualized as a heatmap, where positive values indicate responder-enriched niches and negative values indicate non-responder-enriched niches.

TCR contig annotations were imported from Cell Ranger V(D)J outputs using scirpy. Cell barcodes were standardized by appending sample identifiers, allowing direct matching of TCR and gene-expression profiles, which were integrated into a MuData object using muon/mudata. Immune receptor chains were indexed and quality-controlled using scirpy^13^, retaining cells with at least one non-orphan receptor configuration. Clonotypes were defined using scirpy.tl.define_clonotypes with receptor_arms = "all" and subsequently used to quantify clonal distributions across samples, timepoints and clinical groups. Convergence was defined using scirpy by comparing exact clonotypes with broader amino-acid similarity-based clonotype clusters. CDR3 amino-acid sequences were clustered using the TCRdist metric with a distance cutoff of 15, considering both TCR receptor arms. A clonotype was considered convergent when distinct fine-level clonotypes mapped to the same TCRdist-defined amino-acid clonotype cluster.

### Spatial transcriptomics

Based on hypotheses generated from the single-cell sequencing analyses, we used the 10x Genomics Xenium In Situ platform with a fully custom 480-gene panel, including 18 probes targeting TIE clonotypes (**Supplementary Table 1**) to spatially interrogate the cellular states and interactions of interest. Xenium profiling was subsequently performed on remaining available tumor samples from the POLAR cohort that were not consumed by other experiments, enabling independent spatial validation and extension of single-cell–derived hypotheses.

Spatial transcriptomic profiling was performed on tumor samples from 36 patients enrolled in the POLAR study, yielding a total of 53 Xenium samples across four timepoints: diagnosis (T0, n = 16), screening prior to maintenance therapy (T1, n = 7), during treatment (T2, n = 9), at disease progression (T3, n = 16, including 2 from the same patient), and after progression (T4, n = 5). Samples were obtained from metastatic sites including liver (n = 33), pancreas (n = 7, including 3 fine-needle biopsies), peritoneum (n = 4), and other anatomical locations (n = 9).

Cell segmentation was performed using Proseg, a probabilistic segmentation framework optimized for Xenium in situ transcriptomics data^35^. Proseg integrates transcript spatial density with morphological information to infer cell boundaries. Segmentation outputs, including cell masks and transcript-to-cell assignments, were used to generate cell-level count matrices and spatial coordinates. Xenium outputs, including cell-level transcript counts, spatial coordinates, and morphology-derived cell boundaries, were processed using Scanpy and SpatialData-compatible data structures.

Cell-level quality control metrics included total transcript counts and the number of detected genes per cell. Cells were excluded if they had fewer than 5 detected genes or fewer than 5 total transcripts. To remove segmentation artifacts and merged cell profiles, cells with total transcript counts above the 98th percentile of the dataset were also excluded. Genes were retained if detected in at least one cell. After quality control, approximately 3.2 million cells were retained for downstream spatial analyses.

Cell-level expression matrices were normalized by library size and log-transformed. A k-nearest neighbor graph was constructed from the PCA representation, followed by UMAP for dimensionality reduction. Clustering was performed using PhenoGraph with Leiden community detection. Cell clustering was performed using graph-based community detection (Leiden algorithm), using a high-resolution setting to enable fine-grained identification of spatially distinct cell states. Major tumor, immune, stromal, endothelial, and neural populations were annotated using established marker genes included in the panel. Cell-type annotations were iteratively refined by integrating transcriptional signatures with spatial localization patterns and tissue context. Annotations of morphologically identifiable populations, including epithelial, endothelial, and Schwann cell compartments, were further validated by expert pathological review of matched post-Xenium H&E sections.

To investigate spatial organization and immune exclusion, spatial relationships between cell populations were quantified using cell centroid coordinates derived from Xenium segmentation. For each cell, Euclidean distances to neighboring cells were computed, and neighborhood graphs were constructed using predefined distance thresholds. Cell-type proximity, enrichment, and depletion within peritumoral niches were quantified relative to randomized spatial permutations that preserved overall cell density and tissue geometry. These analyses enabled computation of spatial metrics such as T cell–to–fibroblast ratios within defined peritumoral radii and assessment of immune accessibility to tumor nests. TIE clonotypes were considered confidently detected only when expression was higher in cytotoxic T cells relative to non-immune cell types, thereby minimizing false-positive signals.

## Acknowledgments

This work was supported in part by The Olayan Charitable Foundation and The Tow Foundation. Research support was provided by Philippe Foundation, Institut Servier, Ligue Contre le Cancer, Institut Curie, Fondation de France, Fondation Nuovo Soldati, Fondation Monahan.

## Conflicts of Interest

M.H. reports honoraria from Pierre Fabre, AstraZeneca, Amgen, and Viatris; research funding from Servier; and travel support from Astellas. J.D.S. reports prior consulting for Hoplite Healthcare and serves on the medical advisory board of OpenEvidence, both unrelated to this work. M.M. is an inventor on a patent application related to identifying clones by unique trajectories. K.H.Y. reports a relationship with Ipsen Pharma. D.P. serves on the scientific advisory board of insitro. S.A.V. reports prior consulting for Generate Biomedicines and research funding from Bristol Meyers Squibb unrelated to this work. V.B. reports relationships with Genentech, Merck, and AbbVie. N.R. reports research funding from Pfizer. E.M.O. reports institutional research funding from Arcus, Genentech/Roche, BioNTech, Incyte, AstraZeneca, Elicio Therapeutics, Digestive Care, Agenus, Amgen, Revolution Medicines, and Tango Therapeutics; uncompensated consulting or DSMB activities for Arcus, Actithera, Amgen, Astellas, AstraZeneca, Corcept, Pfizer, Agenus, BioNTech, Ipsen, Ikena, Merck, Immuneering, Moma Therapeutics, Novartis, BMS, Revolution Medicines, Regeneron, Systimmune, Silexion, Tango Therapeutics, and Verastem; travel support from BioNTech, Pfizer, and Revolution Medicines; and other relationships with AACR, ASCO, Imedex, Research To Practice, NIH/NCI, Break Through Cancer, TouchIme, PeerView Voice, and PeerView. W.P. reports institutional research funding from Break Through Cancer, Parker Institute for Cancer Immunotherapy, The Society of MSK, Merck, Astellas, Lepu Biopharma, Amgen, and Revolution Medicines, and consulting for Astellas, EXACT Therapeutics, Revolution Medicines, and Innovent Bio. All other authors declare no competing interests.

