## Supplementary Figures for "Neoantigen-reactive CD8^+^ T cell engagement marks exceptional survivors of pancreatic cancer"

### Supplementary Figure 1

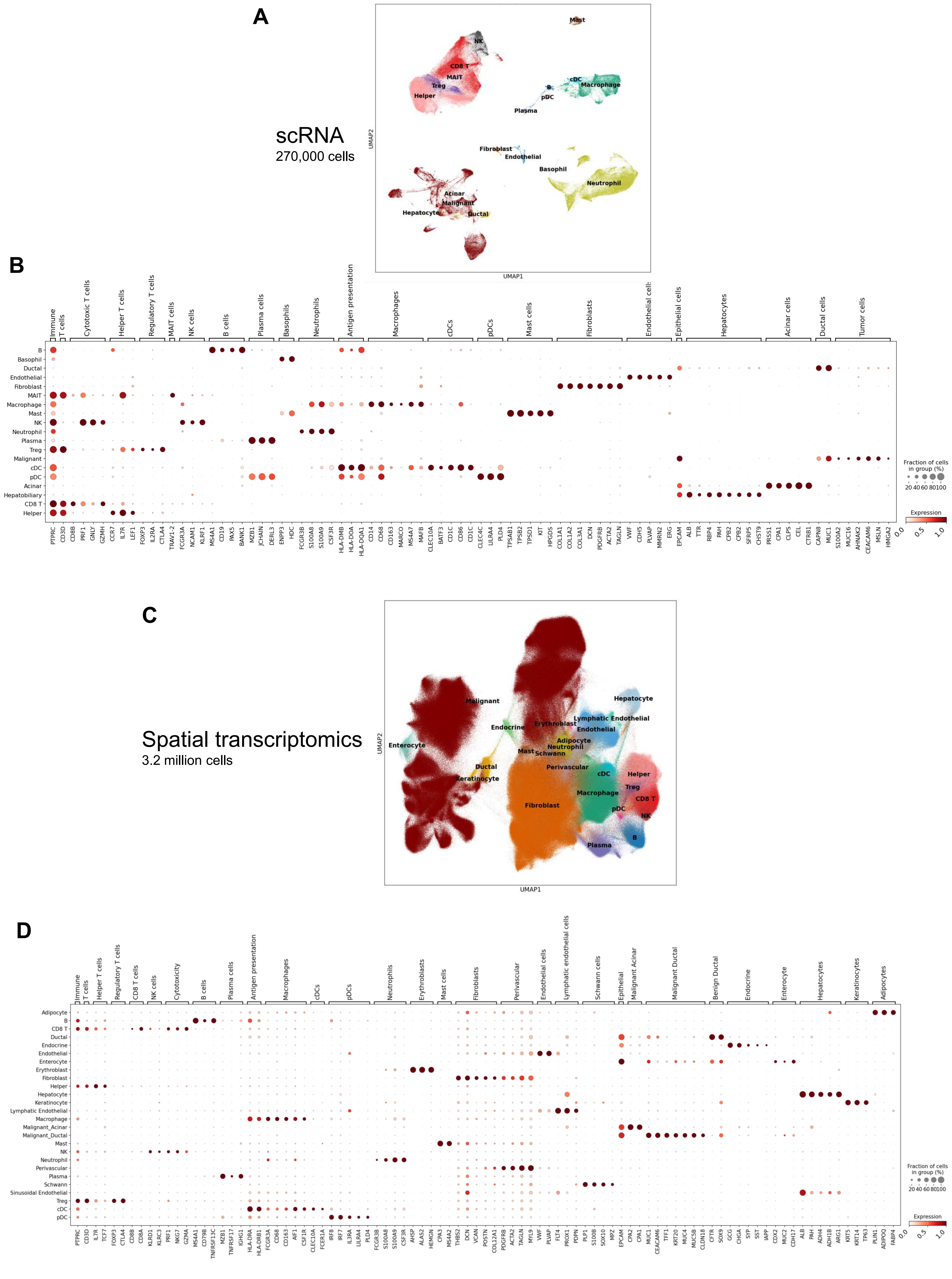

### Supplementary Figure 2

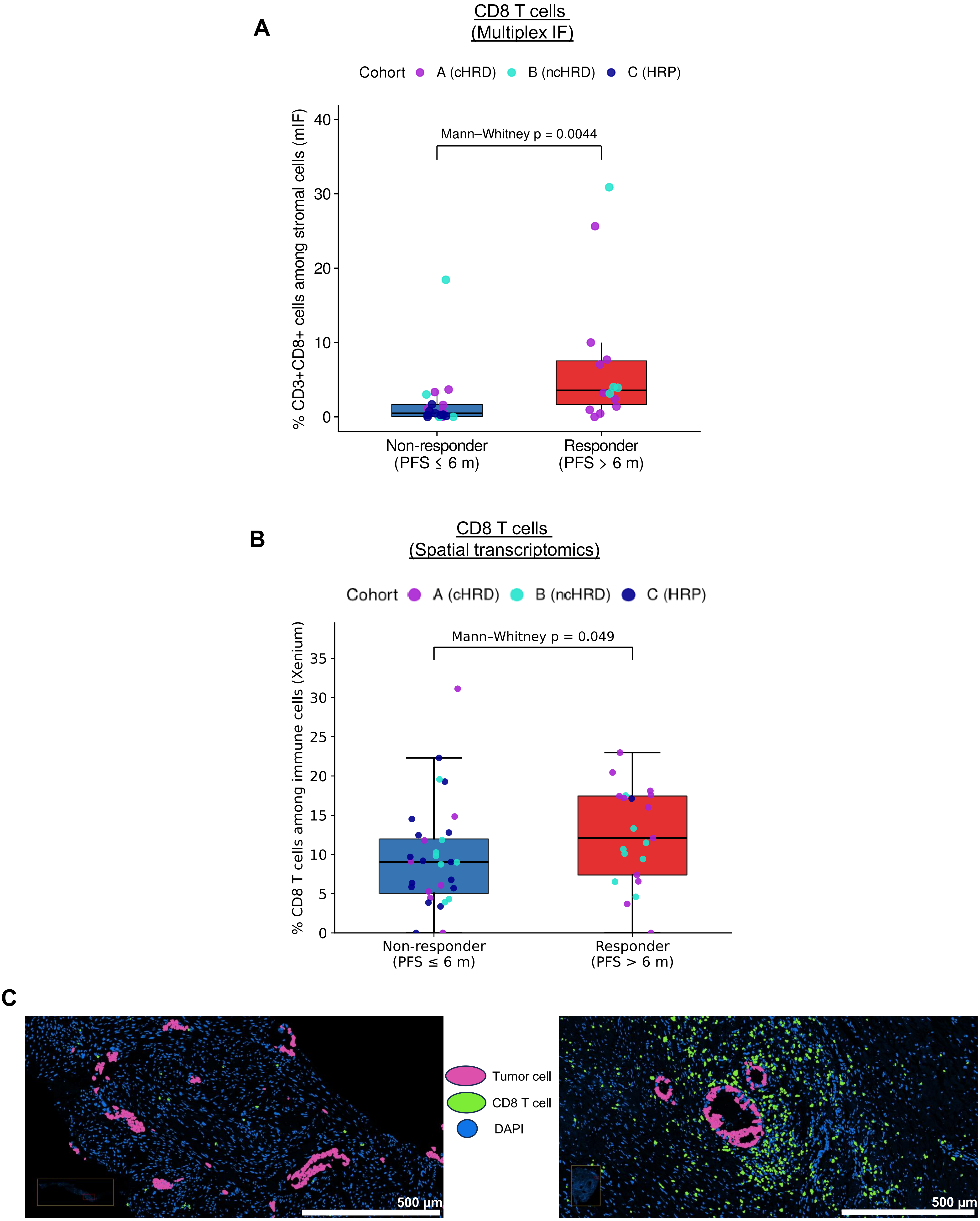


Patient C13, PFS 1.9m
Proportion CD8 T cells among immune cells 5.9%

Patient A04, PFS 46.4m
Proportion CD8 T cells among immune cells 18.1%

**Supplementary Figure 3**

A

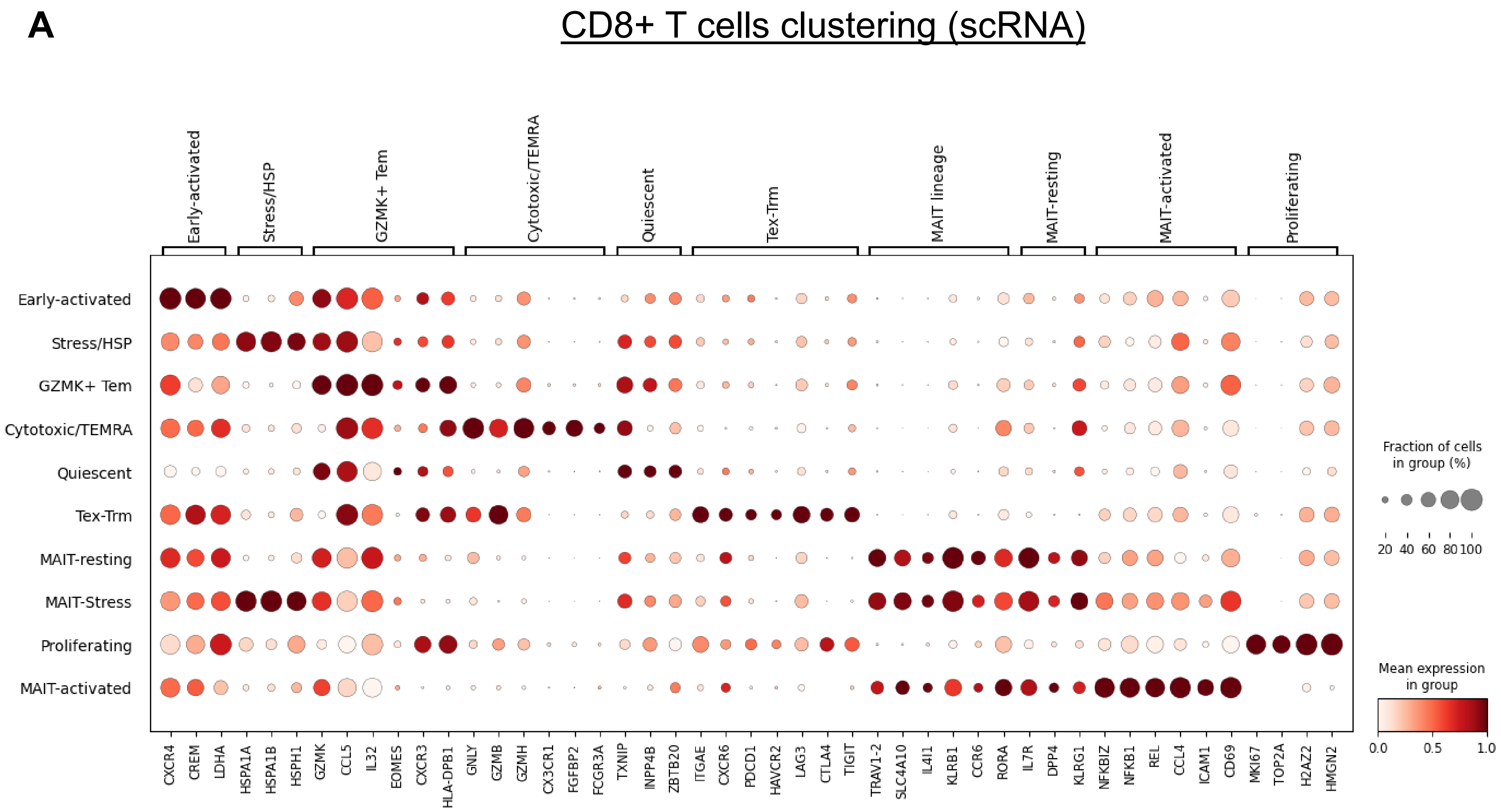


B

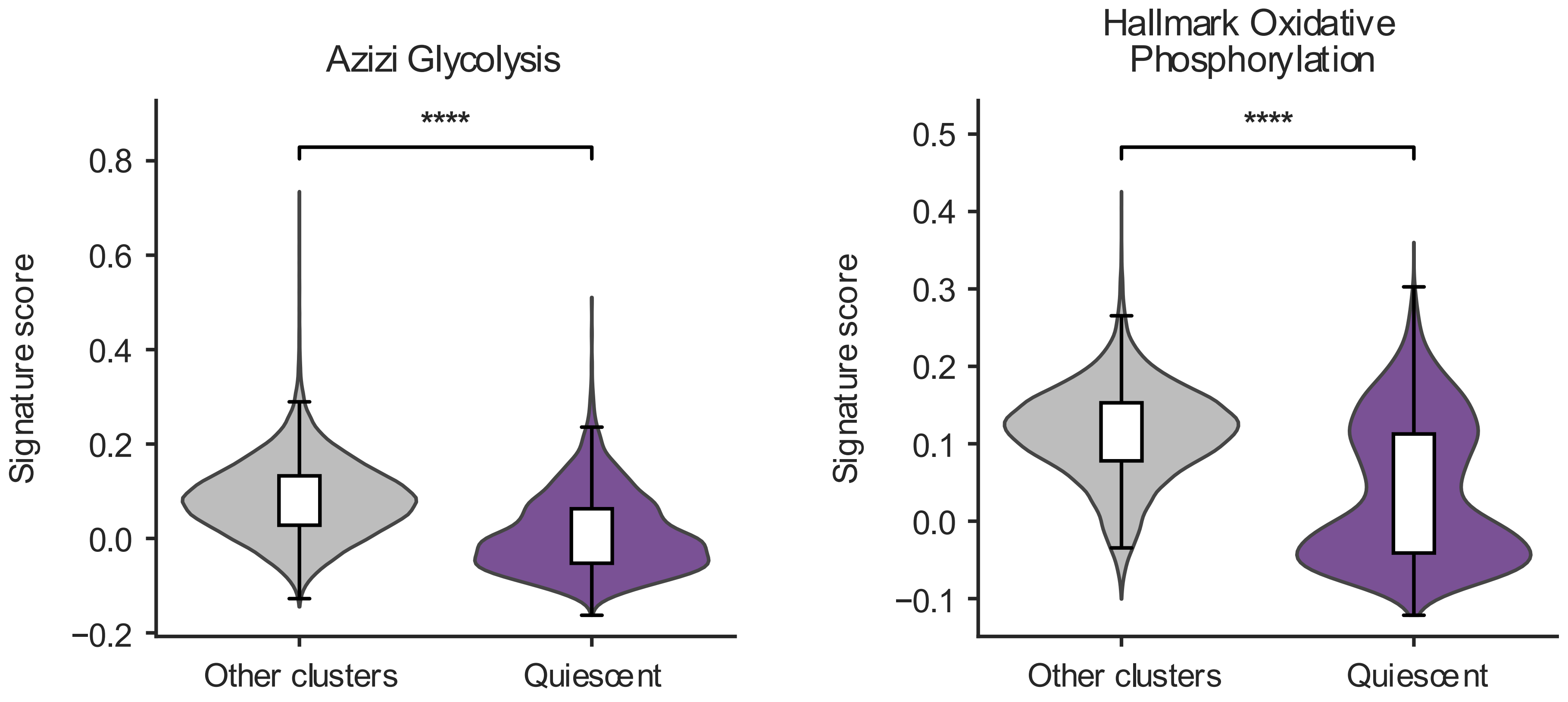

### Supplementary Figure 4

**A** Progression-Free Survival

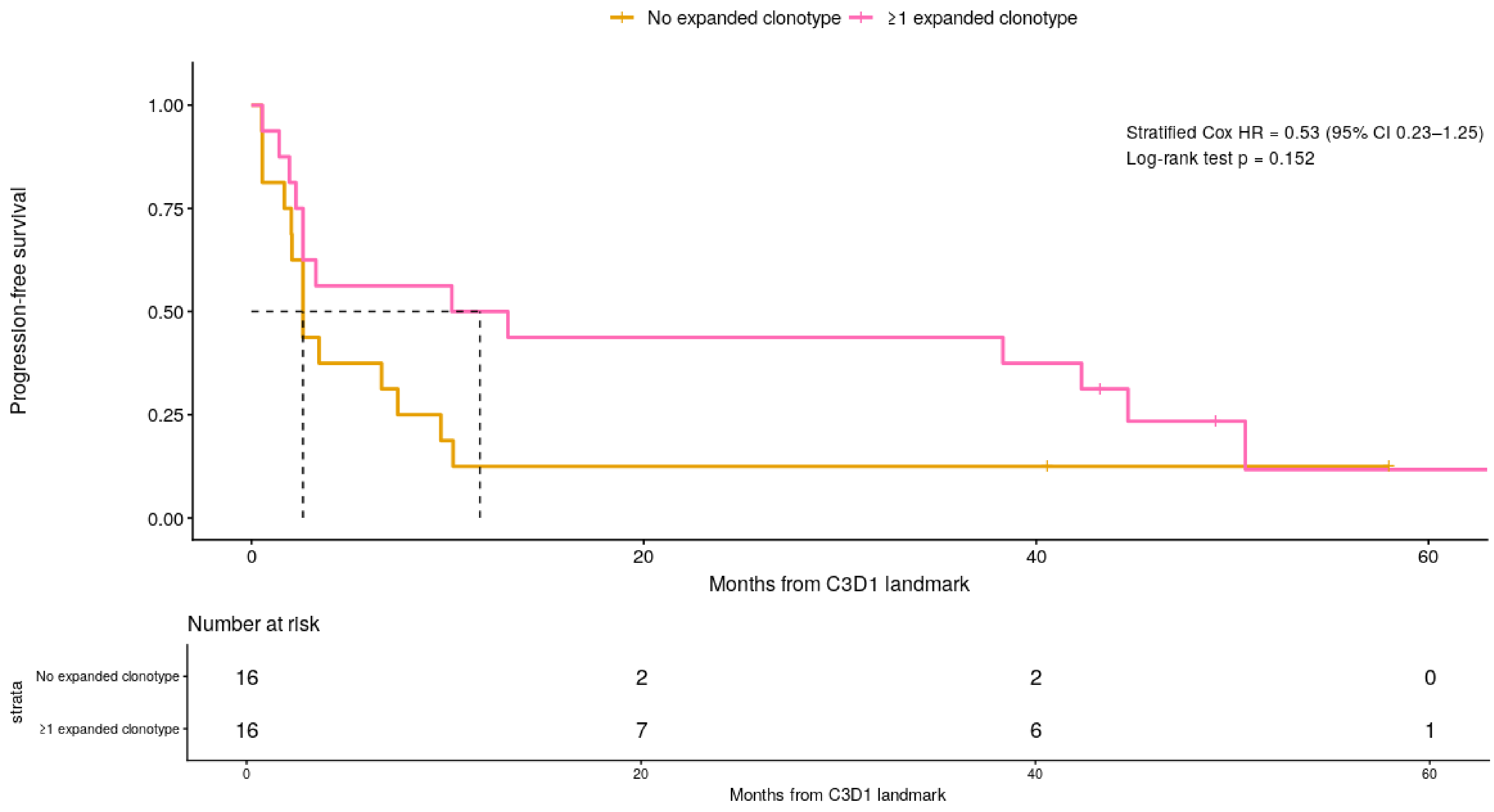


**B** Overall Survival

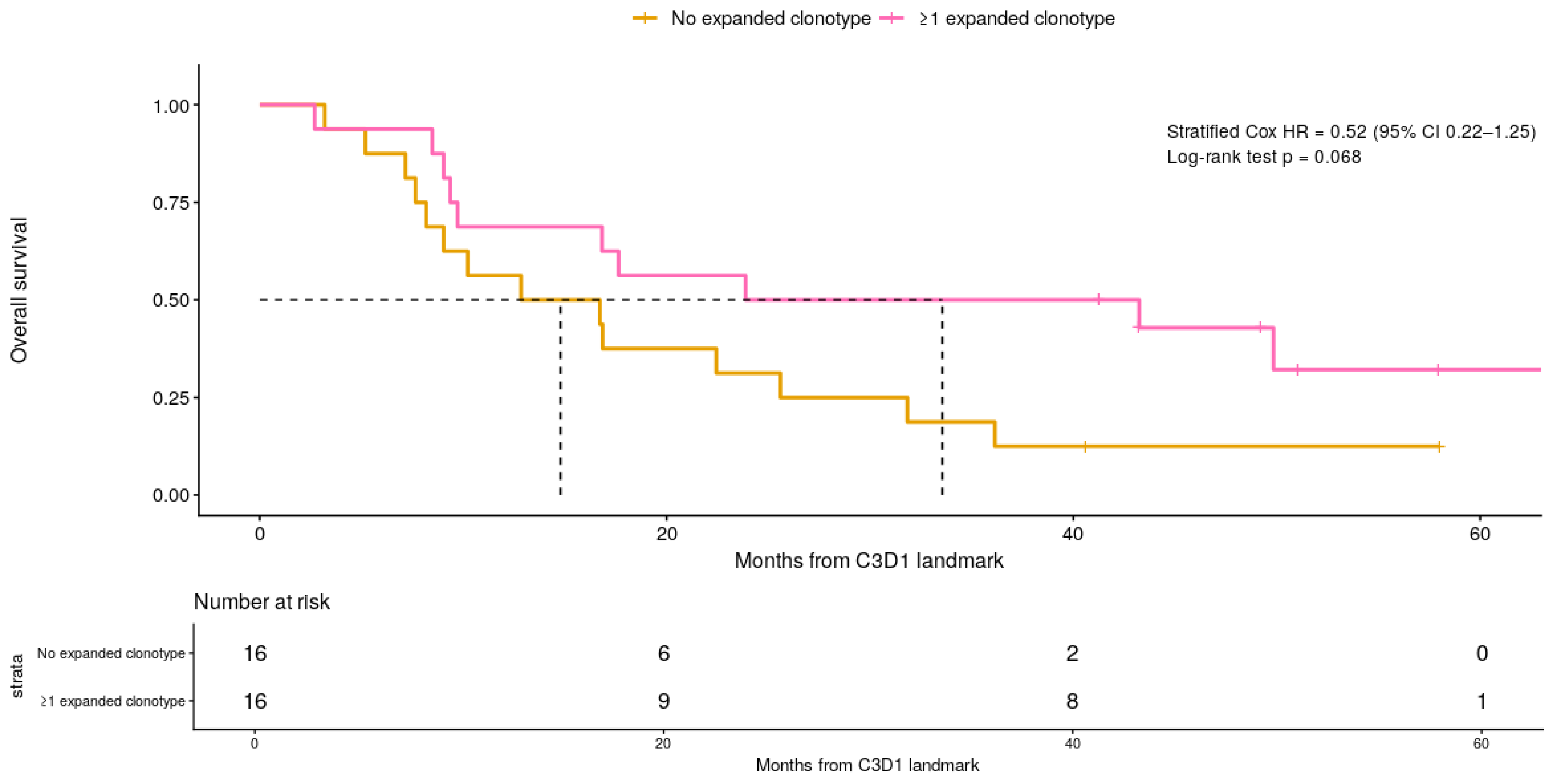

### Supplementary Figure 5

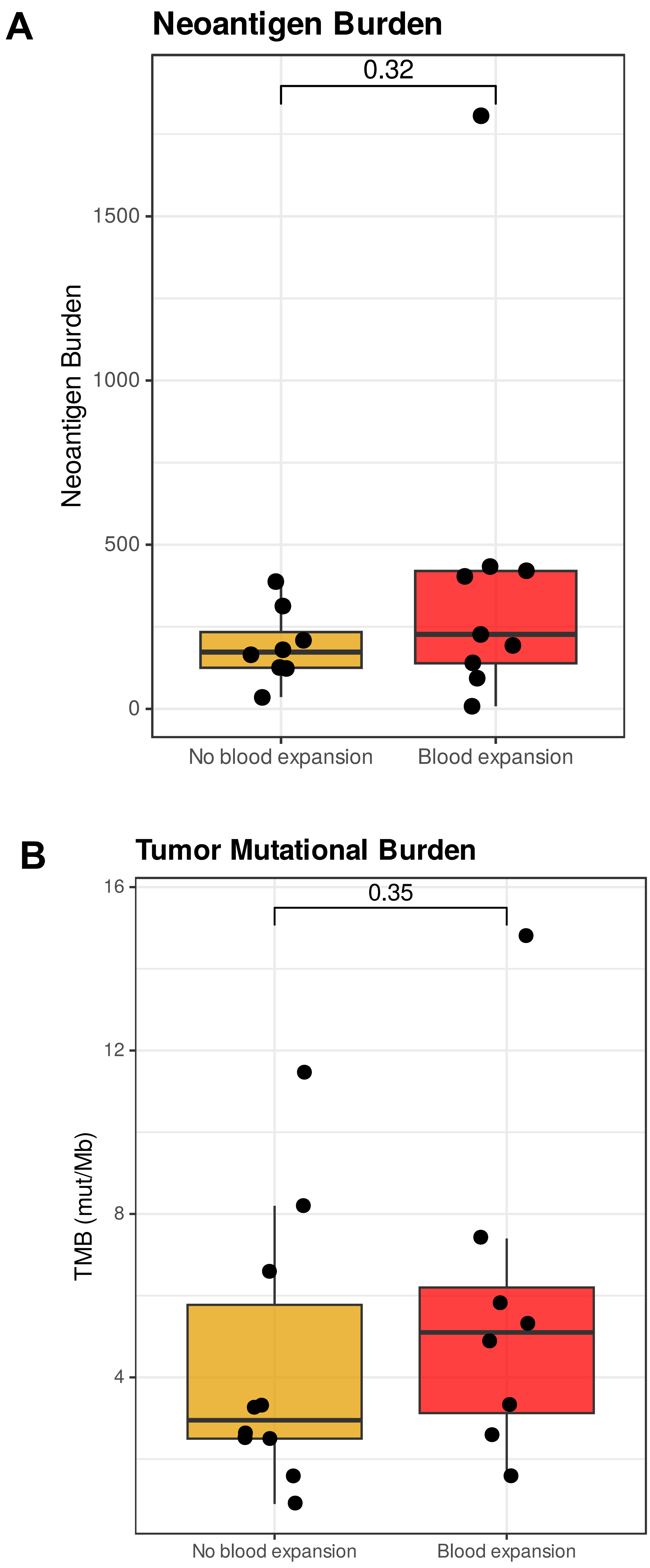

#### Supplementary Figure 6

**A**

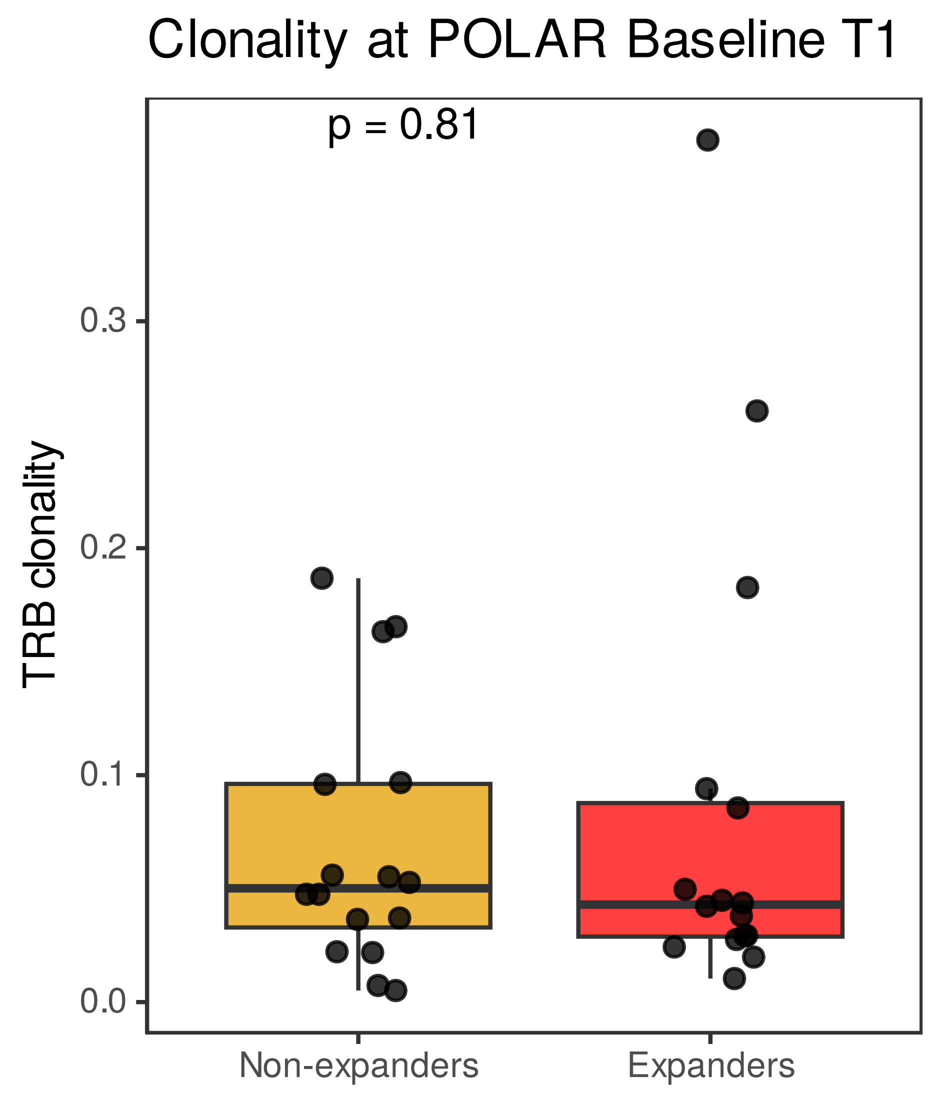


**B**

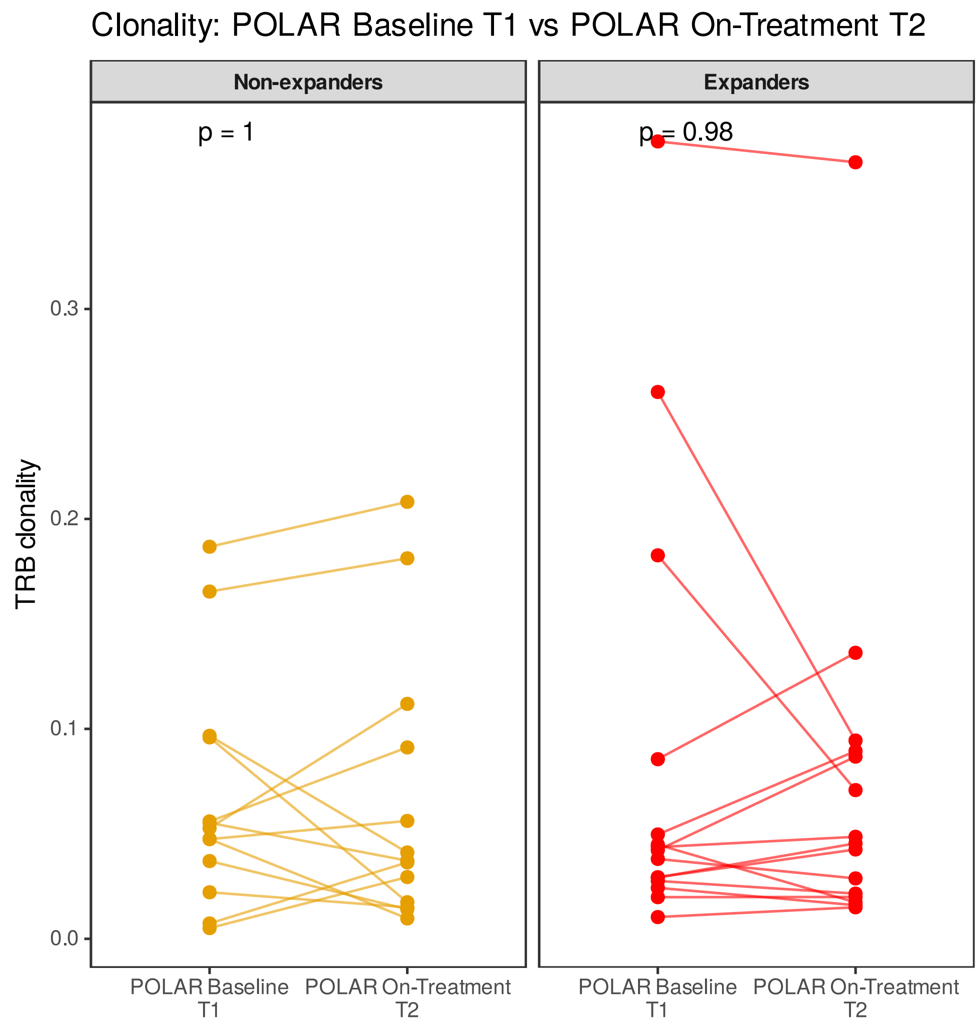

**Supplementary Figure 7**

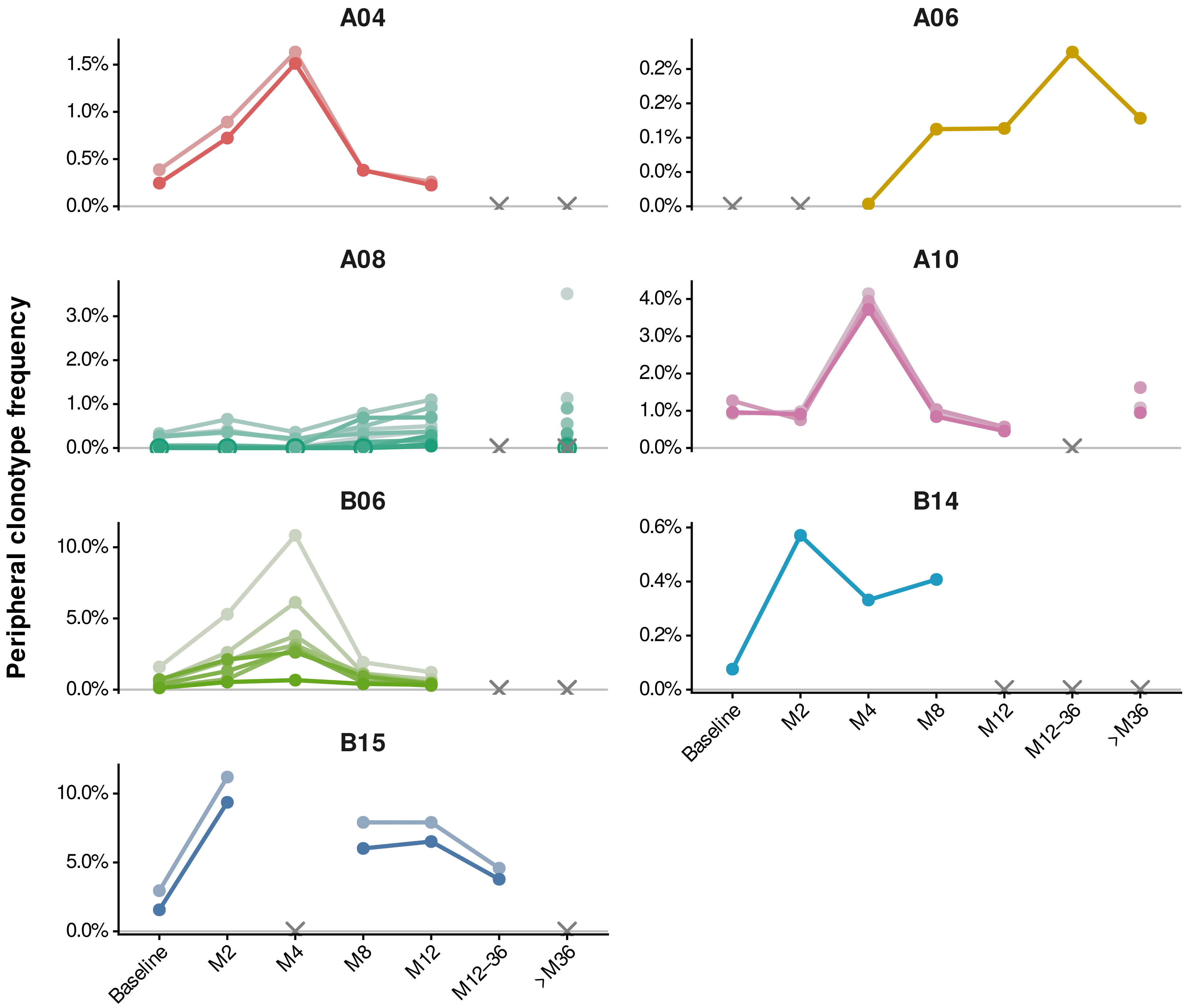

#### Tumor tissue CD8 T cells in TIE+ patients (scRNA)

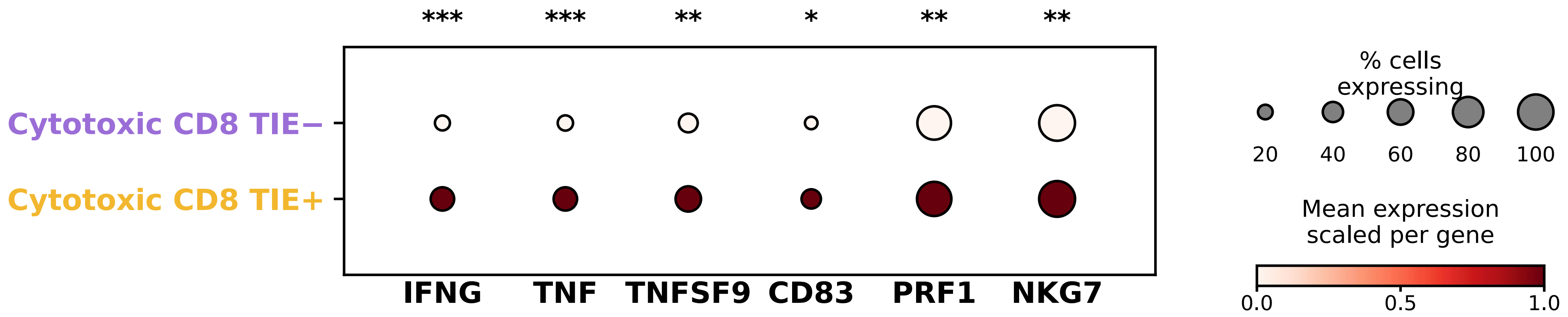

Tumor tissue CD8 T cells in TIE+ patients

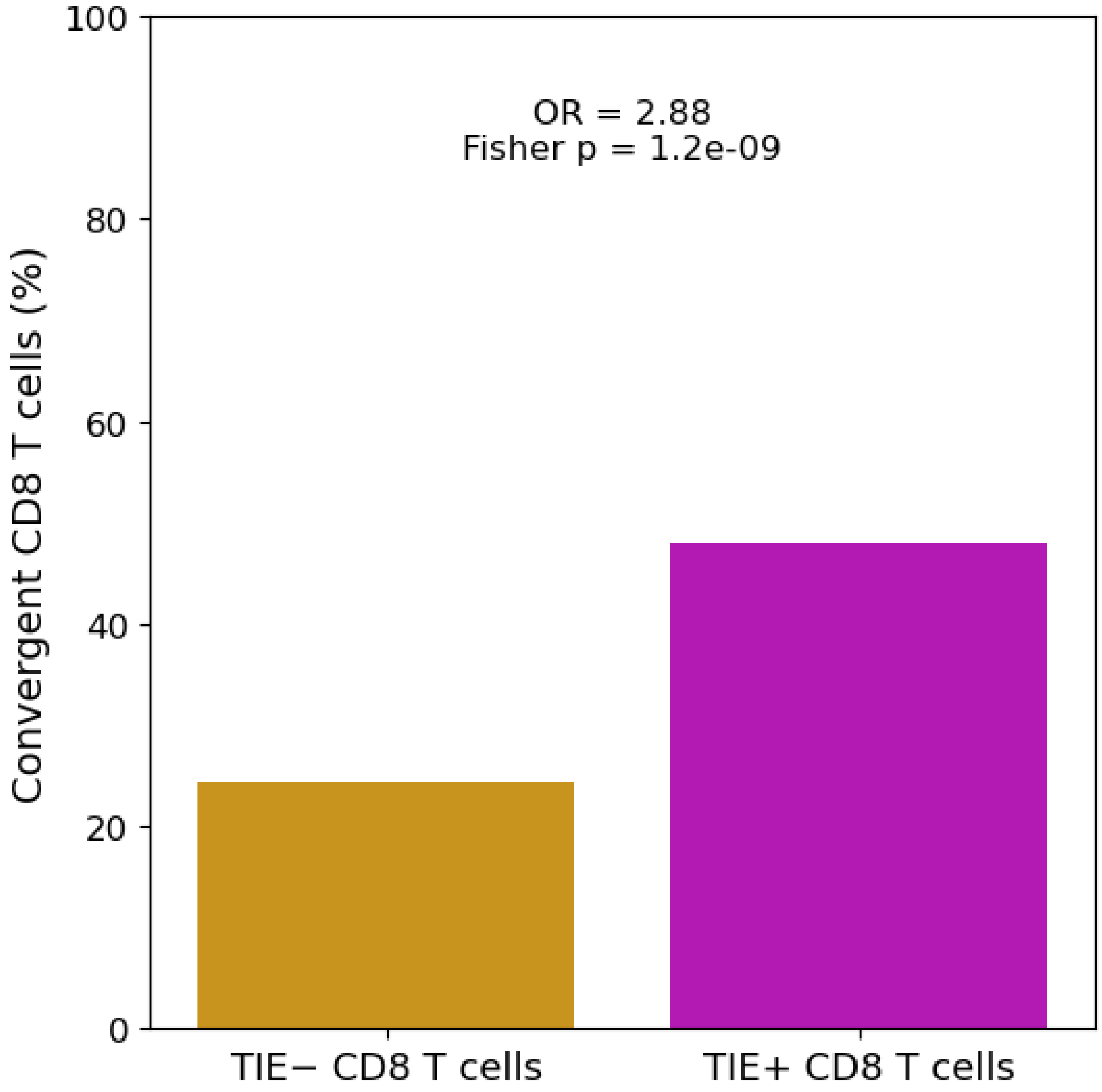

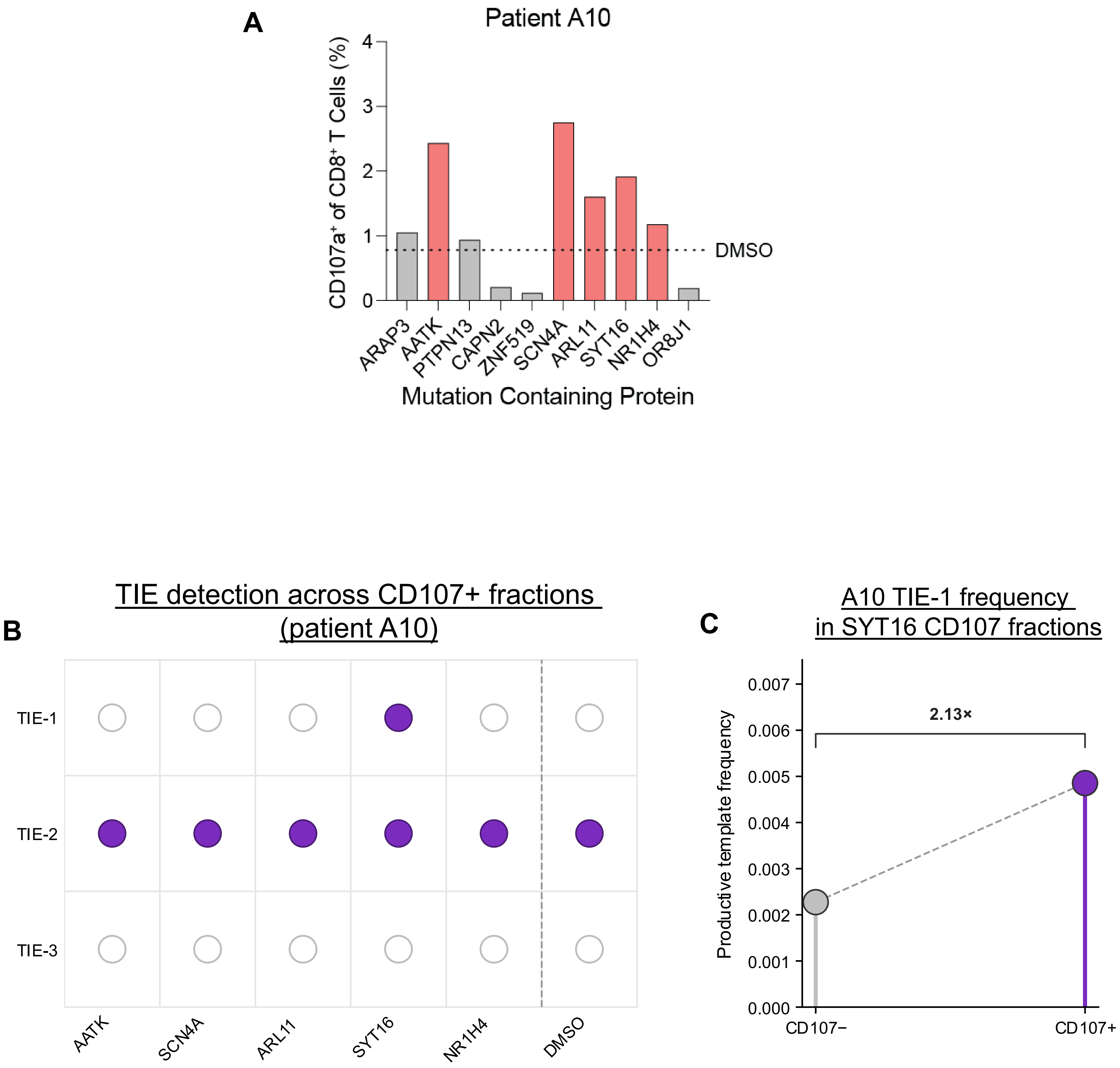
A
Patient A10
CD107a⁺ of CD8⁺ T Cells (%)
DMSO
ARAP3
AATK
PTPN13
CAPN2
ZNF519
SCN4A
ARL11
SYT16
NR1H4
OR8J1
Mutation Containing Protein
B
TIE detection across CD107+ fractions (patient A10)
TIE-1
TIE-2
TIE-3
AATK
SCN4A
ARL11
SYT16
NR1H4
DMSO
C
A10 TIE-1 frequency in SYT16 CD107 fractions
Productive template frequency
2.13×
CD107−
CD107+

### Detection of peptide-source genes in tumor cells at POLAR Baseline

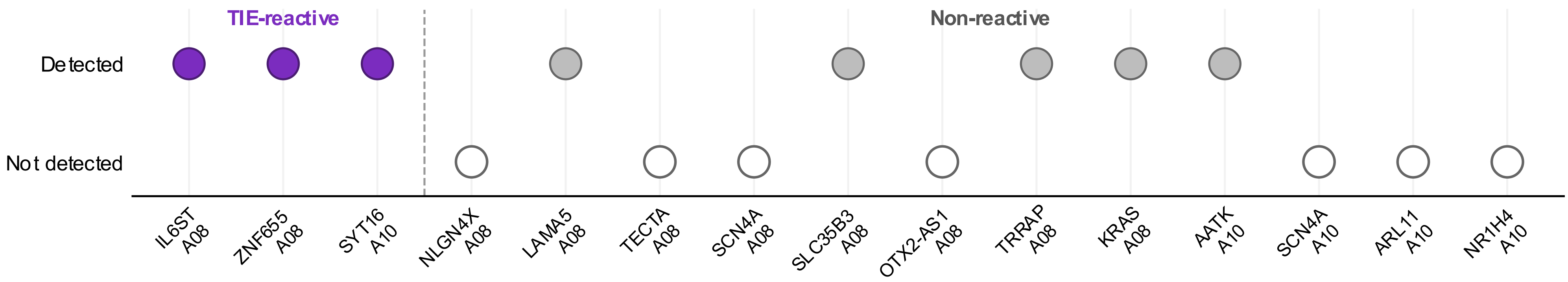
